# Identification of novel loci regulating dormancy in barley and association with hypoxia sensitivity

**DOI:** 10.1101/2024.07.22.604677

**Authors:** Lochlen Farquharson, Bahram Samanfar, Raja Khanal, Wubishet Bekele, Elizabeth K. Brauer

## Abstract

Low seed dormancy is an essential trait in malting barley since malting involves rapid and uniform induction of germination. At least two major QTLs on chromosome 5H, SD1 and SD2, regulate dormancy in multiple barley populations, and additional genetic regions are thought to be involved. To identify novel genetic loci that might be associated with dormancy, a panel of genotypes from diverse agro-ecosystems were evaluated alongside two Canadian biparental populations for germination rate. Association mapping revealed QTLs within the SD1 and SD2 loci in the Canadian populations, while neither of these loci were linked to dormancy in the diversity panel. The diversity panel identified 14 additional marker-trait associations, including novel genetic loci. An alanine aminotransferase (*AlaAT1*) underlies dormancy regulation at the SD1 allele and is thought to help mitigate the suppressive effects of hypoxia on respiration during grain fill. Additional testing with four genotypes carrying either dormant or non-dormant *AlaAT1* alleles revealed that dormant alleles had increased hypoxia sensitivity and hypoxia-responsive gene expression during grain fill. Together, this work indicates that multiple genetic regions influence dormancy and suggests that hypoxia influences dormancy establishment in barley.

**Highlight:** Dormancy is regulated by distinct genetic regions in North American barley compared to African barley. The SD1 locus influences dormancy in North American lines and genotypes with the dormant SD1 allele are more sensitive to hypoxia.

## Introduction

Since plants have limited motility, they must regulate their seed germination to ensure the best chances of survival in a dynamic environment (Willis *et al*., 2014). This selectivity is dependent on two interrelated processes: germination and dormancy. Dormancy is the resistance that a seed has to germination when conditions are suitable for germination and is established during seed development. Dormancy is influenced by genetic and environmental factors such as light, temperature, water content, light, and oxygen (Gubler et al., 2005, Koornneef et al., 2002). During the final stages of seed maturation, seeds are desiccated and typically remain dormant until water is imbibed from the external environment, which is the primary signal to begin germination (Chin et al., 1989). Despite metabolism being predominantly halted in the desiccated seed, the dormancy level is not constant and will decrease over time in a process known as after-ripening (Holdsworth et al., 2008). Germination, dormancy, and after-ripening are three biological processes that actively improve the odds that the plant will enter vegetative growth with the best chance of survival.

In the cereals, dormancy is regulated by cross-talk between hormones, reactive oxygen species (ROS) and hypoxia. During seed maturation, ABA accumulation rapidly increases in the embryo and peaks at 35 days after pollination, which corresponds with the highest level of dormancy (Suzuki et al., 2000; Chono et al., 2006). While ABA has an important role in dormancy establishment, its influence is complex and involves interactions with GA. For example, Chono *et al*., 2006 observed higher ABA content corresponded with increased dormancy levels throughout barley grain maturity, while Romagosa *et al*., 2001 found no relationship between dormancy and ABA content in barley grown in different environments. In maize plants, ABA-deficient mutants germinate precociously except when GA was inhibited, suggesting that interactions between GA and ABA influence dormancy release (White et al., 2000). ABA activates changes in genes expression during grain fill by binding the PYR/PYL/RCAR receptor proteins (Tuan et al., 2018). The ABA-receptors form a complex with a protein phosphatase (PP2C), inhibiting its phosphatase activity. This releases PP2C-mediated inhibition of the SnRK2 kinase activity which subsequently turns on transcription factors including the ABRE-binding protein/ABRE-binding factors (ABRE/ABF) inducing expression of ABA-responsive genes (Tuan et al., 2018). GA signaling is repressed by the DELLA protein, which binds the GA and GIBERALLIN INSENSITIVE DWORF 1 (GID1) to form the GA-GID1-DELLA complex that is marked for degradation (Murase et al., 2008).

Removal of DELLA stimulates the production of *GAMYB* transcription factor that promotes germination (Gubler et al., 1995). Barley spikelets also naturally develop hypoxic conditions in the embryo with the glumella acting as a barrier to oxygen diffusion (Lenoir et al., 1986; Hoang et al., 2013). Hypoxia is likely imposed during establishment of dormancy during grain fill due to the programmed cell death of the photosynthetic external cell layers and restriction of oxygen diffusion by seed coverings (Lenoir et al., 1986). Post-imbibition, limited oxygen availability in the embryo can prevent germination depending on dormancy status, with dormant seeds being more sensitive to hypoxia than non-dormant seeds (Bradford et al., 2008). The mechanisms regulating hypoxia response and its influence on dormancy are not well understood though work in barley suggests that ABA sensitivity increases in lower oxygen environments (Benech-Arnold et al., 2006).

In barley, two main groups of varieties have emerged based on the end use for malting or feed. Malting barley varieties have rapid and uniform germination to ensure product consistency and reduce the storage and production times in the malthouse. The reduced dormancy of malting varieties contributes to their vulnerability to pre-harvest sprouting (PHS), where seeds germinate in the field while still attached to the maternal tissue when conditions are wet (Sweeney et al., 2022). Previous work has identified the SD1 and SD2 loci as major effect QTLs associated with seed dormancy in multiple barley populations, highlighting the importance of the underlying alanine aminotransferase (*AlaAT1*, SD1) and MAPK kinase 3 (*MKK3*, SD2) across multiple barley germplasms (Li et al., 2003; Hori et al., 2007; Ullrich et al., 2009; Nakamura et al., 2017). The SD2 locus is associated with low dormancy at maturity, favorable malt quality, and high PHS susceptibility, with several recent studies suggesting that pleiotropy rather than linkage may be causing this association (Sweeney et al., 2022; Rooney et al., 2023). The alanine aminotransferase1 gene (*AlaAT1*) in the SD1 locus influences variation in dormancy at maturity, and its impact on dormancy increases during after-ripening (Romagosa et al., 1999; Vetch et al., 2020). Multiple alleles in the SD2 locus influence dormancy, and the highly nondormant SD2 will epistatically mask the dormancy effects of SD1 (Sweeney et al., 2022). Some combinations of SD1/SD2 alleles confer the germination levels needed for malting while reducing the potential for PHS, suggesting that it is possible to produce lines with both rapid germination and PHS resistance by optimizing genetic regulation of dormancy (Gao et al., 2003; Sweeney et al., 2022).

To further understand genetic regulation of seed dormancy in spring barley, we evaluated germination in three genotyped populations including the LegCI doubled haploid subpopulation, the SynCH recombinant inbred population and lines from the ICARDA 2014 association mapping (AM-14) diversity panel (Choo et al., 1992; Jui et al., 1997; Luckert et al., 2012, Legge et al., 2014, Amezrou et al., 2018).

## Materials & Methods

### Plant materials

We used the three following barley populations for this investigation; (1) the LegCi population containing 119 doubled-haploid (DH) lines derived from a Leger X CI9831 cross (Choo et al., 1992), (2) the SynCH population containing 88 recombinant inbred lines derived from an AAC Synergy x CH1224-1 cross and (3) 215 genotypes of spring barley from the ICARDA 2014 association mapping (AM-14) panel (Amezrou et al., 2018). The AM-14 panel contains genotypes from row type (88 2-row and 127 6-row) and input adaptation (76 high input, 118 low input and 21 landraces) (Supplementary Table S1). Leger (Léger) is a Canadian 6-row feed cultivar while CI9831 is a 2-row accession (Fejer et al., 1984). The LegCi population has been characterized for disease resistance, seed traits and agronomic traits (Luckert et al., 2012; Chamarthi et al., 2014; Choo et al., 2004). AAC Synergy is a Canadian 2-row malting cultivar (Legge et al., 2014) and CH1224-1 (Pedigree: BM0301-325/AC Kings) is an advanced breeding line from the Ottawa Research and Development Centre. The AM-14 mapping panel has been charactered for morphological traits and resistance to barley diseases (Amezrou et al., 2018, 2021; Gyawali et al., 2018, 2021).

### Plant growth conditions

All genotypes were grown together in a greenhouse to physiological maturity, defined as yellow awns, glumella and first internode region to obtain sufficient seed for dormancy experiments. Three plants were grown in each 6.5 inch fiber pot with Promix(75%):black earth (24%):lime(1%). Plants were grown with a 16-hour photoperiod, 20°C day and 15°C night time temperatures. Plants were watered in the morning and afternoon as needed and were supplemented with NPK 20:20:20 once per week for three weeks when shoots reached approximately 6 inches. From week 3 until heading, the treatment was 35:5:10 followed by 15:15:30 after heading. Heads were harvested at physiological maturity, dried for 48 hours at 37 °C and stored at -20 °C to preserve dormancy. To monitor gene expression plants were grown under the same conditions and heads were frozen in liquid nitrogen and stored at -80°C, 15, 25 or 35 days after anthesis when the awns were first visible.

### Dormancy Testing

Dormancy was quantified by monitoring seed germination in Petri dishes following brief sterilization in 30% bleach for 20 minutes, washing four times in sterile Milli-Q water, and submerging seeds in 0.24% Vitaflow-280 fungicide (UPL AgroSolutions Canada, 11423) for 20 minutes. To simultaneously surface sterilize all lines within a population, seeds from each genotype were placed in 15 mL perforated falcon tubes placed in a custom tube rack within a plastic container containing solutions. After fungicide treatment, 20 seeds were immediately placed embryo (dorsal) side up in 100 mm x 15 mm Petri dishes on one circle of 8.5 cm diameter blue blotter paper (Whatman, #3644) with 11 mL of autoclaved Milli-Q water. The plates were sealed with parafilm and incubated at 20°C in darkness. Each genotype was plated in triplicate Petri dishes, and each population was planted on separate days due to space constraints. The ICARDA AM-14 panel was also split into two days to accommodate the number of lines. The AM-14 lines 12, 59, and 104 were included in both experiments to control day-to-day variability. Germination was determined every 24 hours for the first seven days, day 10, and day 14 after imbibition by scoring the number of seeds with 1 mm coleorhiza emergence as germinated.

To evaluate seed responses to hormones, the same sterilization and imbibition methods were used as described above, but 50 μM ABA was added to the water applied to the filter paper. For ABA sensitivity testing seeds were considered germinated if any detectible radicle growth was observed to separate the effects of ABA on germination from effects on post-germination elongation. To test oxygen sensitivity, seeds were treated with either 21% and 5% oxygen environments using a “Hands-In-Bag” (Spilfyter, WWG3NPA3) disposable atmospheric chamber attached to a cylinder of compressed nitrogen gas where gas was distributed into the bag until the desired level of oxygen was attained. The O_2_ percentage was continuously monitored throughout the experiment using the MO-200 oxygen probe (Apogee instruments). Seeds were either intact or dehulled by inserting fine tipped tweezers behind the rachilla and peeling back the section of the hull covering the embryo. Unless otherwise noted all experiments contained 3 replicate plates containing 20 seeds for each genotype and treatment combination and the experiments were replicated in triplicate.

### Genotyping of Populations

The SynCH RIL lines and their parental lines were genotyped using the two enzyme (*PstI-MspI)* GBS protocol similar to (Torkmaneh et al. 2017; Abed et al. 2022). The genetic information of the parental lines and their progenies were filtered out of the vcf file called by the fastGBS pipeline (Torkmanh et al. 2017) using the Morex V1 IBSCv2 reference genome (Mascher *et al*., 2017). The LegCi population was genotyped using 496 DArT markers as described previously (Luckert et al., 2012). The ICARDA set was genotyped by Illumina arrays (9k and 15K) as described (Amezrou et al. 2018).

### Kompetitive Allele Specific PCR

To evaluate the SD1 and SD2 alleles within genotypes of interest, DNA was first extracted from the leaves of two 3-week-old seedlings from Leger, CI9831, AAC Synergy, CH1224-1, H106-311 and H106-374 along with validation genotypes Esma, AAC Connect and AC Metcalfe from Sweeney et al., 2022*b*. Leaf sections (2.5 cm) were ground by hand with a pestle in 1.5 mL tubes and incubated at 65°C for 30 minutes with 500 μL of extraction buffer (0.1 M Tris-HCl pH 7.5, 0.05 M EDTA pH 8.0, 1.25% SDS). Samples were mixed with 250 μL 6M ammonium acetate before centrifuging for 10 minutes at 10,000 rpm and transferring the supernatant to tubes containing 360 μL isopropanol. Sample isopropanol solution was centrifuged for 10 minutes at 10,000 rpm to pellet DNA. The supernatant was discarded, the DNA was washed with 500 μL of 70% EtOH, dried and resuspended with 80 μL of sterile Milli-Q water. DNA concentration was determined by Nanodrop and diluted to 20 ng/μL for KASP PCR using gene-specific primers based on the methods of Sweeney et al., 2022*b* (Supplementary Table S2). Primer mix was made by adding 12 μL of both forward primers, 30 μL of universal reverse primer and 46 μL water. KASP V4.0 2X Master mix (Lucigen, LGC, KBS-1050-102) was used for PCR reactions in FrameStar 96 well skirted PCR plate, low profile, white walls and black frame (4titude, 4ti-0961). KASP results were measured with Spark microplate reader (Tecan) and analyzed using Kluster Caller software (LGC).

### Statistical Analysis and Association Mapping

Comparisons of small-scale germination experiments was performed using ANOVA with post-hoc Tukey’s HSD or using 2-sided Student’s t-tests (p<0.05) as indicated. For the population germination experiments, broad-sense heritability was calculated by taking the fraction of total phenotypic variation attributed to genotypic variation using a mixed linear model accounting for genotype along with position and shelf in the growth chamber. To test for differences in germination percentage associated with ICARDA sub-populations, row type and adaptation, a nested ANOVA was used with trait nested within day and post-hoc Tukey’s HSD.

Marker filtering and genome association were done using Tassel 5 software (Bradbury et al., 2007). Genotypic data of the diversity panel was filtered to remove markers with minor allele frequency <5%. Marker-trait associations were done using a mixed linear model accounting for population structure (PCA) and kinship matrix. Markers exceeding -log(p-value)= 3 were considered suggestive markers. The 2-row and 6-row genotypes were tested separately for association with germination percentage to account for potential differences in dormancy regulation across sub-populations. In the LegCi population, a linkage map was created using 15252 GBS markers and was placed in 1064 bins using MSTmap across seven major linkage groups (Abed et al., 2022). Each bin was tagged by one SNP marker that is used to produce a linkage map containing 963 markers and spanning a length of 1896 cM (Supplementary Fig. S1). Linkage mapping for the LegCi population was performed using MSTMap (Wu et al., 2008). Genotypic data was associated with phenotypes from 119 genotypes from the LegCi population (Abed et al., 2022) using one- and two-dimensional QTL scans Haley-Knott regression and the multi-QTL model was performed using multiple imputation (Broman and Sen, 2009). A 95% significance threshold was determined using a permutation test with 1000 permutations. Proximity of LOD peaks to known dormancy QTLs SD1 and SD2 were done using the Morex V1 IBSCv2 reference genome (Mascher et al., 2017). Preliminary SynCH linkage map was cleaned using rscripts to remove the high frequency of double crossovers (Latta et al., 2019), and constructed using IciMapping (Supplementary Fig. S1)(Meng et al., 2015). QTL mapping for the SynCH population was done using interval mapping with IciMapping software (Meng et al., 2015). A permutation test with 1000 permutations determined the significance threshold.

### Expression analysis

Transcript levels from whole heads of developing spikes were tested for Leger, CI8831A, H106-311 and H106-374 genotypes. Four heads per genotype were harvested 15, 25 and 35 days after anthesis and flash frozen in liquid nitrogen then stored at -80°C. RNA was extracted from 100 μL of tissue ground in liquid nitrogen and placed in 750 μL of Trizol. Supernatant from a 15-minute spin at 12000xg and 4 °C was mixed with 200 μL of chloroform in phase lock tubes (Quantabio, 2302830). Phase lock tubes were spun at 12000 xg for eight minutes and the aqueous phase was added to 400 μL isopropanol and placed at -20 °C for 24 hours. After the appropriate time interval 650 μL of sample was transferred to the pink column from the Qiagen Rneasy Plant Mini kit (Qiagen, 74904) and spun for 15 seconds at 8000 xg. RNA samples were washed with 700 μL RW1 buffer and two washes with 500 μL RPE buffer. Finally, 50 μL of RNase-free water was added to each column and spun for one minute at 8000 xg to harvest RNA. The RNA was then DNase-treated with 5 μL of TURBO DNase buffer and 1 μL of TURBO DNase enzyme per sample. Samples were incubated at 37°C for 30 minutes followed by inactivation of the DNase enzyme by addition of 1 μL EDTA and incubation at 75°C for 10 minutes. Quality and concentration of RNA was determined using a Nanodrop. cDNA was produced using High-Capacity cDNA Reverse Transcriptase Kit (Applied biosystems, 4368813) according to the manufacturer’s instructions. Quantitative PCR was performed using Power up SYBR master mix in a 10 μL reaction including 1 μL of cDNA and gene-specific primers (Supplementary Table S3) analyzed using a Quant studio 3 (Applied Biosystems). Thermocycler conditions for the qPCR were 2 minutes at 50°C, 2 minutes at 95°C, 40 cycles of 1 second at 95°C and 30 seconds at 60°C. Transcript abundance was normalized to expression of actin and cyclophilin and expressed relative to the Leger 15 days after anthesis (DAA) sample. Experiments were repeated three times with similar results.

## Results

Dormancy was evaluated in three genotyped populations by monitoring radicle emergence over a span of two weeks. The AM-14 panel displayed diverse germination levels approaching a normal distribution at 14 days after imbibition (DAI) with an average germination of 54% (Fig. 1A, Supplementary Data 1). The SynCH and LegCi biparental populations germinated faster in general than AM-14 with the SynCH population requiring less than 24 hours for the majority of the lines to germinate and the LegCi population requiring 4 DAI (Fig. 1A). These results were consistent with the faster germination of the SynCH parental lines compared to the LegCi parental lines (Fig. 1B). Broad sense heritability for germination percentage was high for all three populations including AM-14 (0.67 on 14 DAI to 0.84 on 2 DAI), SynCH (0.67 for 14 DAI to 0.94 for 2 DAI) and LegCi (0.82 for 14 DAI to 0.97 for 1 DAI)(Supplementary Data 1). No significant differences were found when comparing 2-row and 6-row genotypes from AM-14 panel though high input genotypes germinated more than low input genotypes from 3-5 DAI (Supplementary Fig. S2).

**Figure 1.**
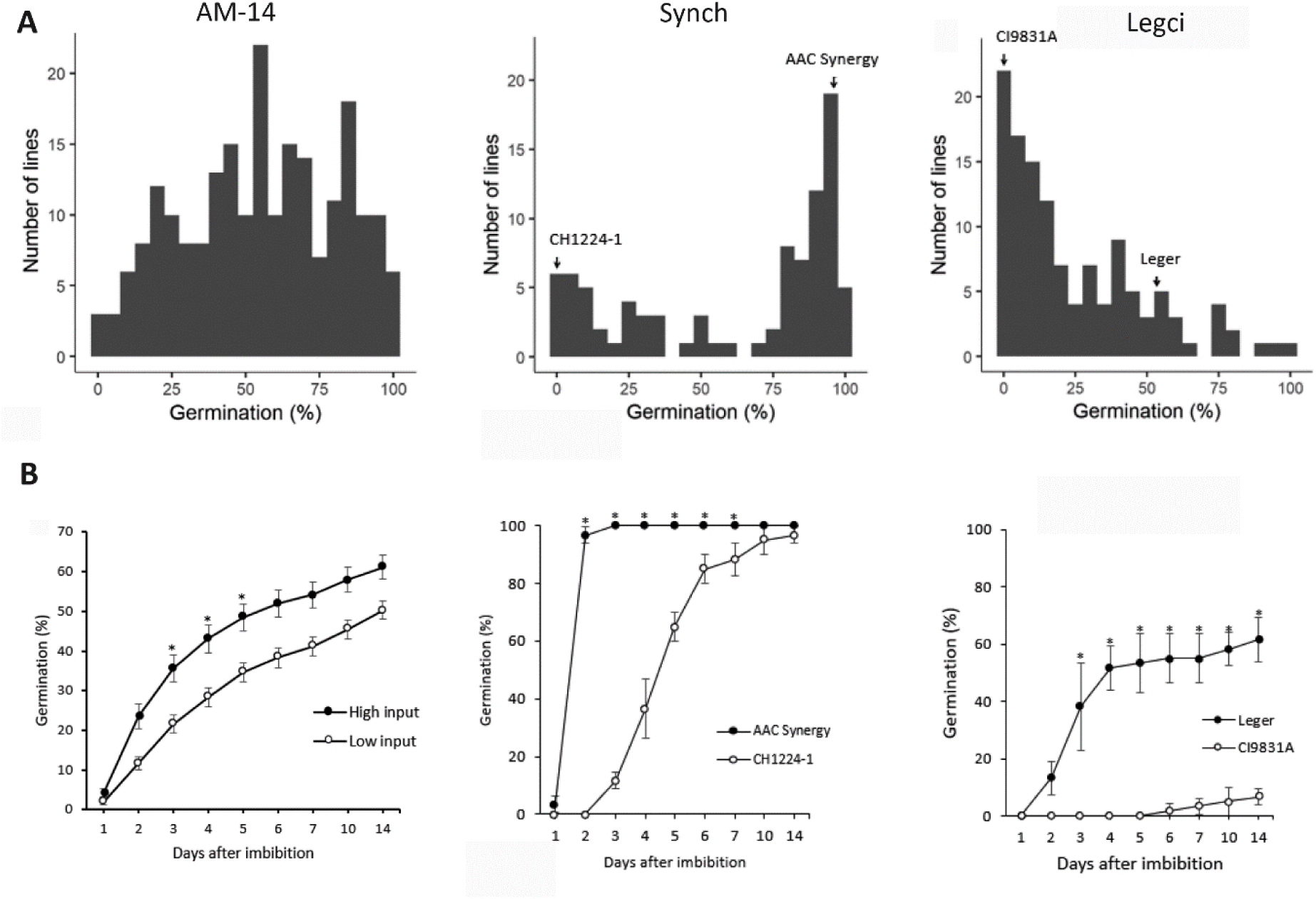
Germination percentage trends for the three surveyed populations including ICARDA AM-14, SynCH and LegCi. For each genotype, 20 seeds were distributed in each of three replicate plates and germination was monitored at the indicated days after imbibition (DAI). (A) The number of lines which display particular germination levels at 14 DAI (AM-14), 2 DAI (SynCH) or 5 DAI (LegCi) is indicated. (B) The average germination percentage ± standard error for all timepoints separated by documented adaptation (118 low input, 76 high input) for the AM-14 panel or for the parental lines is indicated. The * indicates statistically significant difference (p<0.05) from nested ANOVA and the experiment was repeated once.

Previous work has identified the SD1 and SD2 loci as major effect QTLs associated with seed dormancy in multiple barley populations (Li et al., 2003; Hori et al., 2007; Ullrich et al., 2009; Nakamura et al., 2017). To determine if the SD1 and SD2 loci could be contributing to the observed dormancy or lack thereof in our biparental populations, the KASP method was used to distinguish dormant and non-dormant associated SNPs (Table 1). One SNP was found to influence dormancy in SD1 (L214F) by Sato *et al*., 2016, and two in the SD2 locus (E165Q, R350G) where SD1 can be designated as dormant (D) or non-dormant (ND) while SD2 is D, ND or highly non-dormant (ND*), masking any dormancy at the SD1 allele (Vetch et al., 2020). Sweeney *et al*., 2022*b* did not directly test R350G but used a 50k matrix SNP highly correlated with R350G in their population, termed JHI-367342-KASP. As reported previously, AAC Synergy contains two non-dormant SD2 and a dormant SD1 allele (Vetch et al., 2020; Sweeney et al., 2022). The CH1224-1 is dormant for both SD2 alleles and non-dormant for SD1, contrasting the non-dormant parent AAC Synergy at every tested locus. Leger had the dormant allele present at both SD2 loci but was non-dormant at SD1 while CI9831 was dormant or non-dormant for the SD2 allele and dormant at SD1 in line with previous reports indicating the two genotypes of the CIho9831 male parent (Luckert et al., 2012). We also tested two progeny lines H106-311 and H106-374 from the LegCi population as the H106-311 genotype shared 78% of the 963 molecular markers with CI9831 but has the non-dormant allele at SD1 while the H106-374 shared 76% of its markers with Leger but had the dormant allele at the SD1 locus (Table 1, Supplementary Fig. S3). These lines provide an opportunity to compare the effects of SD1 and genes in the immediately surrounding regions for their effect on dormancy as the lines share molecular markers in chromosome 5 surrounding the SD2 region. In line with SD1 having an influence on these lines, the H106-374 (dormant SD1) germinated slower than Leger (t-test: p = 0.004) and H106-311 (non-dormant SD1) germinated faster compared CI9831 (t-test: p= 0.006)(Table 1).

**Table 1:** SNP identity at known seed dormancy QTLs SD1 and SD2 in parental lines for SynCH and LegCi populations and recombinant lines H106-311 and H106-374. Three to six replicate plants were analyzed separately to determine consistency of the result. Germination results are average germination percentage ± standard deviation for 20 seeds germinated in three different plates from a single germination assay used to produce mapping phenotypes. Dormant (D)/ non-dormant (N) refers to G/C (MKK3_E165Q), A/G (JHI-367342-KASP) or G/C (AlaAT_L214F) respectively. The dormant allele is represented by D and non-dormant by N. For the SD2 allele N* indicates both SNPs have the nondormant allele.

| Genotype | Population | SD1 | SD2 | GP 3 DAI | GP 5 DAI | GP 14 DAI |
| --- | --- | --- | --- | --- | --- | --- |
| AAC Synergy | SynCH (parent) | D | N* | 100 ± 0.0 | 100 ± 0.0 | 100 ± 0.0 |
| CH1224-1 | SynCH (parent) | N | D | 12 ± 2.8 | 65 ± 5.0 | 97 ± 2.8 |
| Leger | LegCi (parent) | N | D | 38 ± 15.2 | 53 ± 10.4 | 62 ± 7.6 |
| CI9831 | LegCi (parent) | D | N/D<sup>+</sup> | 0 ± 0.0 | 0 ± 0.0 | 7 ± 2.8 |
| H106-311 | LegCi | N | D | 12 ± 7.6 | 15 ± 5.0 | 25 ± 13.2 |
| H106-374 | LegCi | D | D | 7 ± 5.7 | 15 ± 5.0 | 57 ± 10.4 |

+ mixed results between replicate plants.

To identify the genetic loci that were associated with germination in the 3 different populations, both GWAS and QTL linkage mapping approaches were used. For the AM-14 panel, GWAS was used for germination percentage at 2 DAI and 14 DAI to assess early and long-term germination trends. The 2 DAI data revealed two marker-trait associations (MTAs) on chromosomes 4H (BOPA1_4919-1051) and 7H (BOPA1_4644-1363) with a third on 7H (BOPA2_12_11477) (Fig. 2A, Table 2). The 14 DAI data did not reveal any significant MTAs (Fig. 2B). Separating the AM-14 by 2-row or 6-row lines revealed significant marker associations on chromosome 3H and 7H for the 2-row lines and on chromosome 4H and 7H for 6-row lines (Supplementary Fig. S4). When high input-adapted lines were mapped independently one MTA was found on 7H at 2 DAI and another on 5H at 14 DAI. Low input-adapted lines had six MTAs found across chromosomes 2H, 3H, 6H and 7H. Interestingly, the SD1 locus was not identified as significant in this population as associated with dormancy. One MTA was found on 5H closer to the SD2 locus when high input lines were mapped independently at 14 DAI, an unusual timepoint for SD2 association since non-dormant SD2 is typically highly non-dormant but no markers were mapped to 5H for the earlier timepoint (Table 2, Supplementary Fig. S4)(Sweeney et al., 2022).

**Figure 2.**
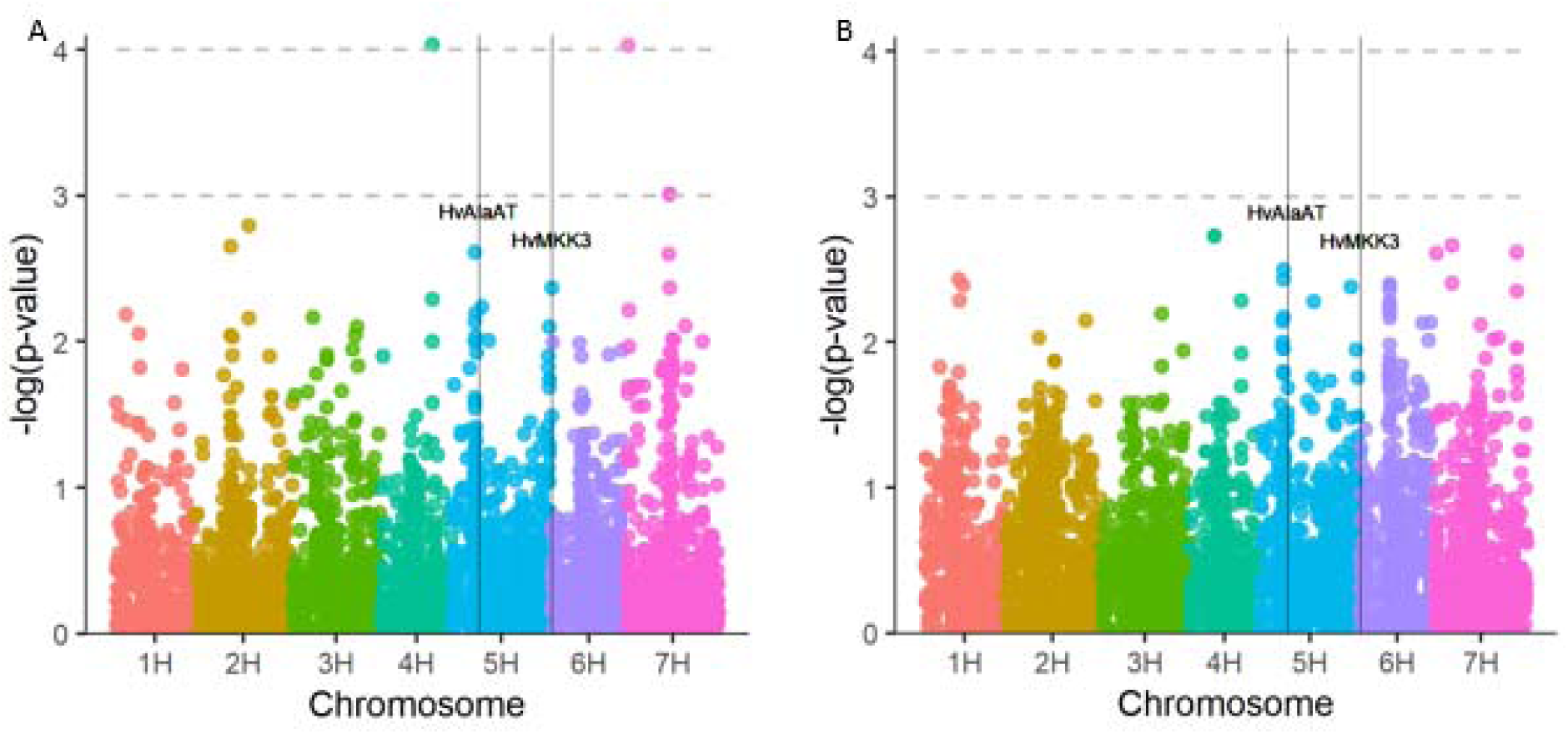
Association between (A) 2 DAI or (B) 14 DAI germination percentage and 4689 SNP-based markers for the AM-14 panel. Association was determined with a mixed linear model accounting for kinship structure using Tassel 5.0 software. Markers located with -log(p-value) ≥ 3 were considered significant. Approximate location of SD1 (HvAlaAT) and SD2 (HvMKK3) was done by locating the closest marker to each position respectively on the Morex v3 reference genome.

**Table 2:** Marker-trait associations for germination percentage with the ICARDA AM-14 population with at 2 or 14 days after imbibition (DAI). Association mapping was performed independently on: all 215 lines, 88 2-row, 127 6-row, 76 high input or 118 low input genotypes. MTA was considered significant at -log(p-value) ≥3. Information about markers was obtained from T3/barley and Morex v3 physical position was determine by BLAST using Grain Genes. The * indicates markers identified in multiple association maps.

| Marker | DAI | Row type | Input Adaptation<sup>1</sup> | Chromosome | $-\log(p\text{-value})$ | Position (cM)<sup>722</sup> |
| --- | --- | --- | --- | --- | --- | --- |
| **BOPA1_4919-1051\*** | 2 | 2/6 | High / Low | 4H | 4.04 | 584593042 |
| **BOPA1_4644-1363\*** | 2 | 2/6 | High / Low | 7H | 4.03 | 2832220 |
| BOPA2_12_11477 | 2 | 2/6 | High / Low | 7H | 3.01 | 248325635 |
| SCRI_RS_67208 | 2 | 2 | High / Low | 3H | 3.3 | 2536489 |
| BOPA2_12_30399 | 2 | 2 | High / Low | 3H | 3.1 | 488628723 |
| **BOPA1_4919-1051\*** | 2 | 6 | High / Low | 4H | 3.4 | 584593042 |
| SCRI_RS_4520 | 2 | 6 | High / Low | 7H | 3.02 | 605413584 |
| BOPA2_12_31173 | 14 | 2 | High / Low | 7H | 3.5 | 9155301 |
| **BOPA1_4644-1363\*** | 2 | 2/6 | High | 7H | 3.1 | 2832220 |
| SCRI_RS_234527 | 14 | 2/6 | High | 5H | 3.4 | 571092964 |
| SCRI_RS_736 | 2 | 2/6 | Low | 6H | 3.8 | 521284380 |
| SCRI_RS_13615 | 2 | 2/6 | Low | 7H | 3.3 | 7360724 |
| BOPA1_3965-353 | 2 | 2/6 | Low | 3H | 3.1 | 564661364 |
| BOPA1_3359-1118 | 2 | 2/6 | Low | 7H | 3.1 | 6871605 |
| SCRI_RS_136586 | 2 | 2/6 | Low | 7H | 3.09 | 572855259 |
| SCRI_RS_235860 | 2 | 2/6 | Low | 2H | 3.07 | 567601498 |

To determine if similar genetic regions underlie germination rate in the biparental populations, we conducted QTL mapping. In the SynCH population, QTL mapping was done using 2827 markers (Supplementary Fig. S1) on germination percentage at 2 DAI since this phenotype had the greatest difference between parental lines. Germination percentage at 2 DAI (H^2^ = 0.94) is significantly associated with markers in the telomeric region of 5H (Fig. 3A, Table 3). One of the MTA for 2 DAI in SynCH was at marker chr5H_659541058 which explained 10% of the variation in 2 DAI germination percentage (Table 3). The marker chr5H_659541058 is located 11.6 Mbp away from *MKK3* (SD2) based on the Morex V1 IBSC v2 reference genome (Mascher et al., 2017). There were 174 genetic markers that could not be assigned to a particular linkage group for the SynCH population and were mapped as a separate linkage group. Three markers from this group were strongly associated with 2 DAI germination percentage, together explaining 51% of the variation (Fig. 3B, Table 3). In the LegCi population, QTL mapping based on 963 markers (Supplementary Fig. S1) and germination percentage from all timepoints revealed consistent MTA trends with the strongest association occurring 5 DAI (H^2^ = 0.86). A multiple imputation based multi-QTL model explained 46% of the variation in germination percentage at 5 DAI and indicated two significant loci for association with germination percentage on 2H and 5H, respectively (Fig. 3C, Table 3). The 2H loci had a minor effect on germination percentage, explaining 3.3% of the variation, while the 5H loci had a major effect on germination percentage, explaining 39.6%. The QTL on 5H was located around marker chr5H:490954074, which has an allele frequency of roughly 54% Leger and 45% CI9831 identity, respectively. When comparing the germination percentage of genotypes with the dormant versus non-dormant SNP at chr5H:490954074 which is the closest marker to SD1 1.88 Mbp away, the dormant genotypes predominantly clustered under 20% germination at 5 DAI while nondormant genotypes are more evenly distributed between 0-80% germination categories (Fig. 3D). This distribution suggests the dormant allele may be maintaining dormancy rather than the non-dormant allele breaking dormancy since non-dormant lines are not necessarily pushed to high germination. This result is in agreement with our KASP analysis demonstrating variation in dormancy-associated SNPs in SD1 between the LegCi parental lines but not in the SD2 locus (Table 1). Overall, we identified 14 unique genetic regions underlying dormancy from the AM-14 population, a single locus that is proximal to SD2 in the SynCH population, and two loci in the LegCi population, one of which is proximal to SD1. Interestingly, we did not identify strong control by SD1 and SD2 in the AM-14 population, indicating that genetic regulation of germination differs in genetic backgrounds from the ICARDA collection compared to the Canadian genotypes. In the Canadian germplasm, SD1 and SD2 were identified as major effect QTL controlling seed dormancy, with SD1 in the feed variety and SD2 in the malt variety.

**Figure 3.**
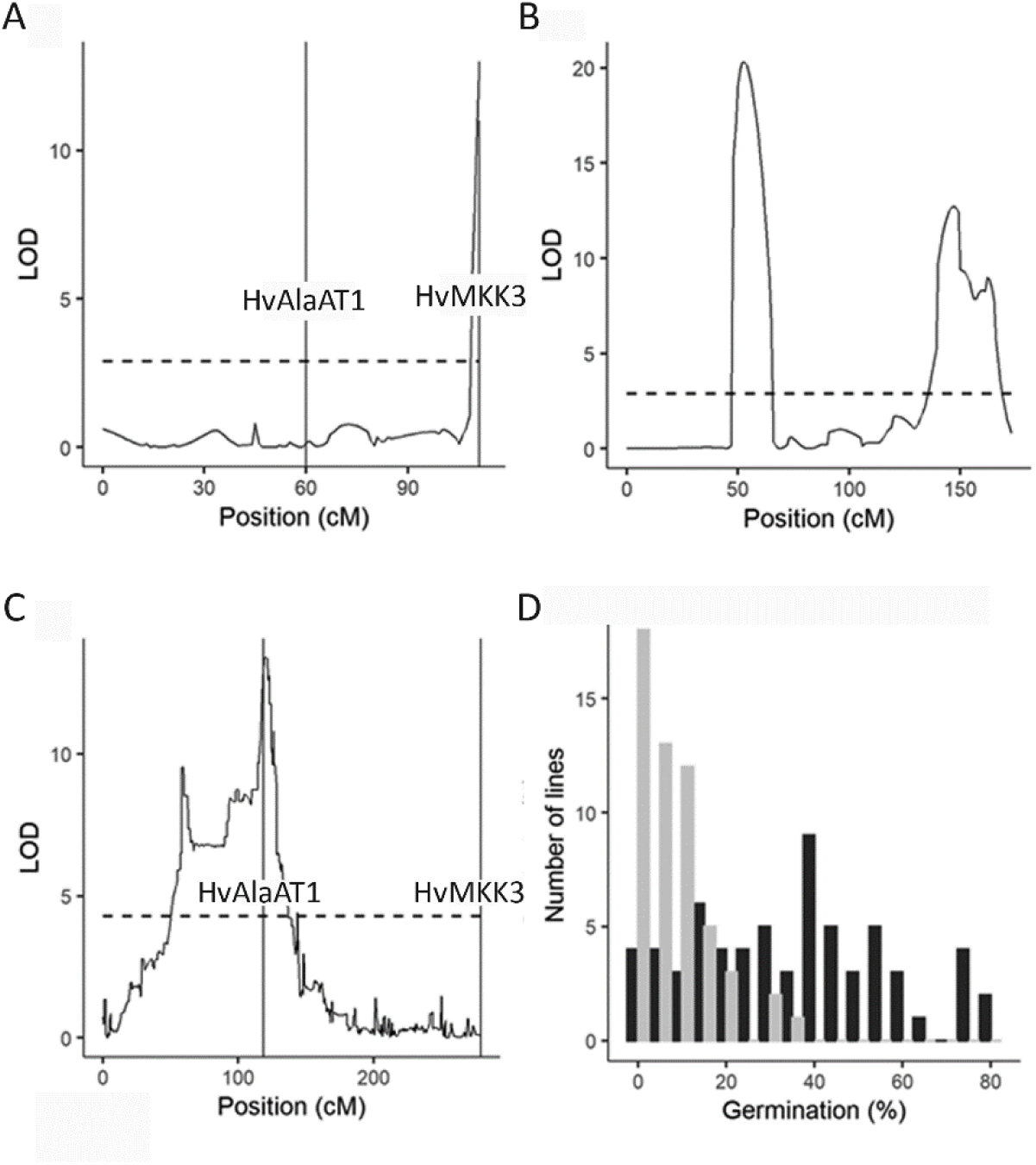
Association of germination percentage with markers from the SynCH and LegCi populations. (A) Association of germination percentage 2 DAI with 2827 markers from the SynCH population revealed a QTL on chromosome 5H and a number of unassigned markers for linkage group 8 (B). Significance threshold is 2.9 and was calculated via 1000 permutations. (C) For the LegCi population (119 lines) at 5 DAI, association of 963 genetic markers revealed a QTL on chromosome 5H. The LOD was estimated with Haley-Knott regression where the significance threshold is 4.28 determined with 1000 permutations. Approximate location of SD1 (HvAlaAT1) and SD2 (HvMKK3) was determined by identifying the marker with the closest physical position using the Morex v1 IBSCv2 reference genome. Logarithm of odds was calculated using interval mapping with the IciMapping software (Meng et al., 2015). (D) The distribution of germination percentage in the LegCi population 5 DAI separated by identity at marker chr5H:490954074 to either Leger (non-dormant: black) or CI9831 (dormant: grey).

**Table 3:** Marker-trait associations identified in the LegCi and SynCH populations. Association was estimated in the LegCi population using Haley-Knott regression (r/qtl; Broman and Sen, 2009) and in the SynCH population using standard interval mapping (IciMapping; Meng et al., 2015). Association genotypic markers was performed with germination percentage (GP) from multiple days with the strongest association for each population presented.

| Population | Trait | Chromosome | Marker | Percent variation explained (%) |
| --- | --- | --- | --- | --- |
| Legci | 5 DA\| GP | 2H | chr2H_632650121 | 3.3 |
| Legci | 5 DA\| GP | 5H | chr5H_490954074 | 39 |
| Synch | 2 DA\| GP | 5H | chr5H_659541058 | 10.26 |
| Synch | 2 DA\| GP | Unassigned | SUN_92613782 | 17.29 |
| Synch | 2 DA\| GP | Unassigned | SUN_247107242 | 17.2 |
| Synch | 2 DA\| GP | Unassigned | SUN_249641186 | 16.9 |

The LegCi population identified SD1 as the only major effect QTL controlling dormancy in this population indicating that the underlying *HvAlaAT1* significantly impacted germination. The mechanisms underlying the influence of *HvAlaAT1* on dormancy are not well understood though expression occurs during the grain maturation phase between anthesis and physiological maturity when dormancy is established (Chono et al., 2006; Sato et al., 2016). In Leger, the *HvAlaAT1,* but not four other predicted alanine aminotransferases, were induced during grain fill where expression of *HvAlaAT1* increased 75 times at 15 days after anthesis compared to at anthesis (Supplementary Fig. S5). To determine how ABA, GA and hypoxia-responsive genes are regulated in barley with dormant versus non-dormant alleles of SD1, we tested expression at 15 days after anthesis (DAA) in Leger, CI9831 and progeny (H106-374, H106-311) lines. At 15 DAA, *HvAlaAT1* expression was similar between lines, as was the expression of other alanine aminotransferases (Supplementary Table S4). Genes responsive to GA (*GA2ox3, GA3ox2*) and ABA (*NCED1*, *ABA8H-1, ABI5*) were also similar across lines (Supplemental Table S4). This trend corresponded with a similar level of germination inhibition response to ABA application across the lines (Supplementary Fig. S6). However, a hypoxic stress-responsive gene in the ethanol fermentation pathway encoding an alcohol dehydrogenase (ADH) was significantly higher in the lines carrying the dormant SD1 allele including CI9831 and H106-374 (Fig. 4)(Bui et al., 2019). Barley embryos are typically exposed to hypoxic conditions (5% oxygen) in mature seeds due to the tightly adhering glumella which limits oxygen diffusion (Lenoir et al., 1986). To determine if hypoxic conditions impacted germination the LegCi parental lines, we monitored germination in 5% oxygen compared to 21% oxygen, representative of non-hypoxic conditions, and observed that hypoxia significantly suppressed germination in all lines (Fig. 5A,B). To compare embryo hypoxia sensitivity in our four lines of interest, we next dehulled the seeds prior to exposing them to hypoxic conditions to ensure the oxygen levels were consistent and were not affected by differences in seed coverings across lines. In dehulled seeds, all genotypes germinated rapidly in 21% oxygen with visible signs of radicle growth within 1 DAI and approximately 100% germination by 2 DAI (Fig. 5C,D, Supplementary Fig. S7). In the 5% oxygen environment, all dehulled genotypes had suppressed germination for the first 3 DAI compared to the 21% oxygen environment. Dehulled seeds from both genotypes featuring the non-dormant allele of SD1 (Leger and H106-311) were less sensitive to the inhibitory effect of 5% oxygen on germination compared the two lines featuring the dormant allele (CI9831 and H106-374) and had significantly higher germination percentage starting at 2 DAI (Fig. 5C, D). Together with our gene expression analysis, this suggests that the non-dormant *HvAlaAT1* is associated with reduced hypoxia sensitivity in barley seeds.

**Figure 4.**
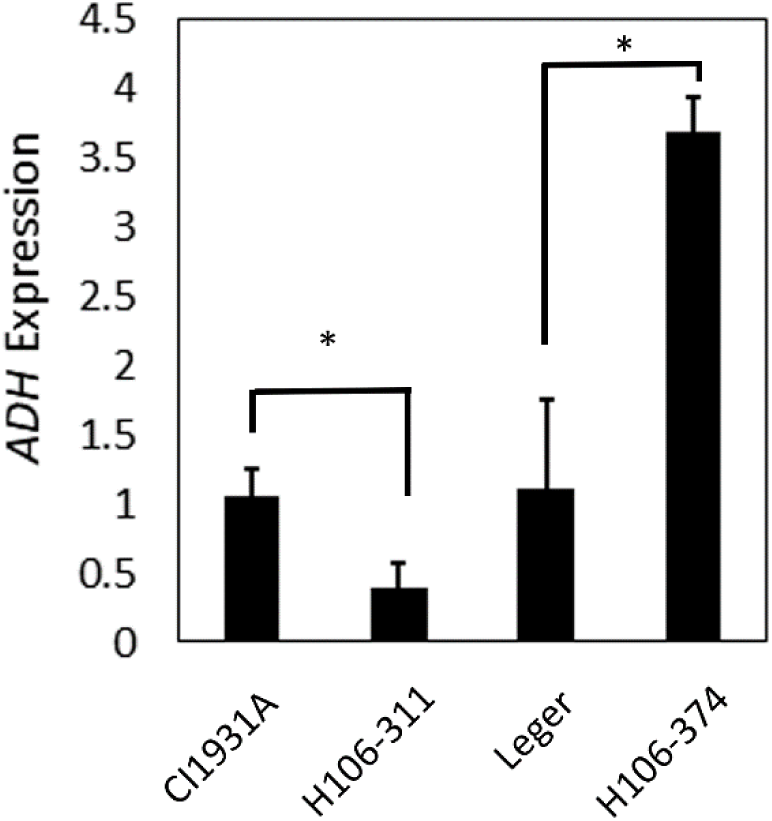
Relative expression of the alcohol dehydrogenase *ADH* gene during grain fill in genotypes segregating for the SD1 allele. Barley spikes were harvested at 15 days after anthesis from 4 separate plants per genotype and tested for gene expression by quantitative RT PCR to compare genotypes carrying the dormant (D, CI9831, H106-374) or non-dormant (ND, Leger, H106-311) SD1 allele. Expression was normalized to actin and cyclophilin expression as outlined in methods and expressed relative to the parental lines (CI9831, Leger). Results were pooled across three independent experiments (n=9-12) where the mean and standard error are shown here. Significant differences were determined by a 2-tailed t-test and indicated by * (p<0.05).

**Figure 5.**
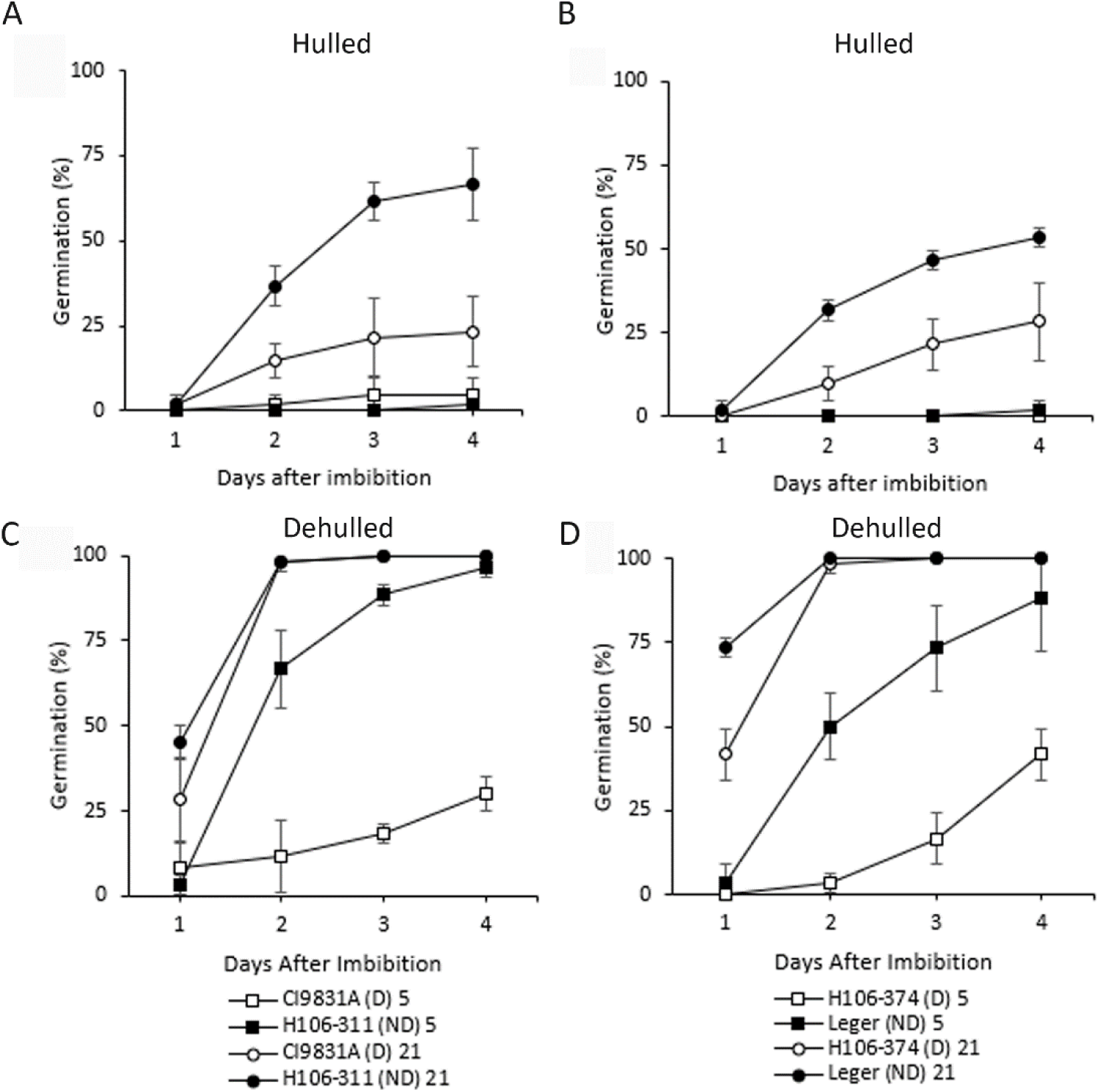
Association between SD1 and hypoxic tolerance during germination of hulled (A, B) and dehulled (C, D) barley caryopsis. Tested lines contain either the non-dormant (ND; closed) or dormant (D; open) allele of HvAlaAT1 with a genetic background similar to CI9831 for A) and C) or Leger B) and D). 20 seeds germinated at room temperature in air (∼21% O_2_; circle) or 5% O_2_ (square) environments. Values represent average germination percentage ±standard deviation (n=3). The experiment was replicated three times for each genotypic pair with consistent findings.

## Discussion

Breeding high-quality malting barley varieties has released the natural dormancy that exists within this species, leading to issues with germination on maternal plants. Ideally, malting barley would have high dormancy prior to harvest and fast, consistent release of dormancy during malting. Understanding the genetic and physiological mechanisms regulating dormancy is essential for balancing the timing of dormancy release in malting barley. Here, we identified 17 novel and known regions of the genome associated with dormancy through association mapping in the AM-14 panel and biparental linkage mapping in the SynCH and LegCi populations of spring barley. The majority of associations were identified with germination percentage 2 DAI indicating the strongest genetic control of germination occurs soon after imbibition. SynCH and LegCi populations each identified strong regulation by SD2 and SD1 respectively, though the SD1 and SD2 alleles were not associated with dormancy in AM-14. This is consistent with previous work demonstrating that the SD2 allele explained 36% of the variation in dormancy in the Canadian malting variety AC Metcalfe which was included in the production of AAC Synergy (Zhang et al., 2012; Legge et al., 2014). AAC Synergy was also recently included as the common parent in a connected half-sib population that identified SD2 linked MTA for regulation in PHS resistance further implicating SD2 as the major controlling factor of dormancy in this genotype (Sweeney et al., 2020). In the LegCi population, a major effect QTL at SD1 explained 39% of the variation in dormancy with Leger contributing the non-dormant allele. The QTL was located over an extended region of chromosome 5H similar to previous findings in Australian genotypes and could indicate multiple genes in this region influencing germination (Hickey et al., 2012). Distribution of lines suggested the dormant SD1 allele is maintaining dormancy rather than the non-dormant allele being involved in promoting germination which agrees with previously published trends (Nagel et al., 2019). Indeed, the effect of SD1 on dormancy in barley has been found to increase with after-ripening time but still has an effect on primary dormant seeds harvested at physiological maturity (Vetch et al., 2020).

The AM-14 panel was sourced from ICARDA in Morrocco which includes genotypes adapted to abiotic stresses and favorable production conditions (Amezrou et al., 2018). Therefore, the reduced average dormancy in high input lines may be due to the prevalence of malting-related genetic backgrounds which are produced to be non-dormant at maturity. This suggests that in malt-focused germplasms minor associations influencing dormancy may be selected out making low input germplasm a potential source of novel dormancy regulation genes. For example, the AM-14 panel revealed an association with the 24-methylenesterol C-methyltransferase 2 (*SMT2*) gene on chromosome 4H that was also found in American germplasm (Correa-Morales, 2013). SMT2 is a key enzyme in the production of the sterols sitosterol and stigmasterol (Rogowska and Szakiel, 2020). Sterols have been observed to influence seed germination in *Arabidopsis* and could also play a role in cell membrane recovery from desiccation by modulating membrane fluidity (Rogowska and Szakiel, 2020; Shimada et al., 2021). In addition, two of the MTAs located on chromosome 7H were identified were also identified for root length 12 days after imbibition in drought stressed barley seeds (Thabet et al., 2018). Additional investigation into the minor effect alleles within the AM-14 would help to clarify the prevalence of dormant and non-dormant alleles in malting varieties and their distribution in North American populations.

While SD1 has been linked to dormancy regulation in multiple populations of barley, the role of the underlying alanine aminotransferase in regulating dormancy is not well understood. Our work indicates that the expression of *HvAlaAT1* is induced during grain fill but the amplitude of its expression is not different between lines carrying dormant versus nondormant SD1 alleles similar to findings by Sato *et al*., 2016. Our work on a small set of lines indicates that *HvAlaAT1*’s role in establishing dormancy seems to be independent of ABA and GA transcriptional regulation or ABA sensitivity. Previous work with mutant barley lacking *HvAlaAT1* expression indicated that mutants had significantly higher ABA content compared to controls suggesting a role for the gene in ABA production or stability (Hisano et al., 2022). Thus, the connection between *HvAlaAT1* and hormonal regulation remains unclear and further work is needed to conclusively establish if alanine aminotransferase influences hormone production or responses.

Our results indicate hypoxia-response genes are induced during grain maturation, with *ADH* being upregulated and differently expressed in SD1 non-dormant lines. The association of *HvAlaAT1* allele with embryo sensitivity to 5% oxygen suggests that *AlaAT1* could influence dormancy from either increased level of oxygen accumulation within the seed or decreased requirement for oxygen due to altered metabolic processes. Alanine aminotransferase is an enzyme that reversibly converts pyruvate and glutamate to alanine and 2-oxogluterate, and is transcriptionally induced in response to hypoxia (Muench et al., 1998; Duff et al., 2012). Studies on oxygen status in pea roots found that the plant’s control of pyruvate levels is critical to limiting oxygen consumption, and therefore limiting hypoxia on internal tissues (Zabalza et al., 2009). Studies on waterlogging in *Lotus japonicus* found an increase in AlaAT activity and accumulation of alanine under waterlogging conditions, implicating AlaAT for a role in controlling pyruvate levels during hypoxia (Rocha et al., 2010). Since alanine accumulates under hypoxia, greater activity of alanine aminotransferase could result in less pyruvate for respiration and greater oxygen retained in the tissue. It is not clear how after-ripening occurs at the molecular level but the process is associated with lipid oxidation and ROS accumulation nonenzymatically in the desiccated seed (Oracz et al., 2007; Bouteau et al., 2007). Lipid oxidation is a self-accelerating reaction which generates ROS and the partial pressure of oxygen can greatly affect the eventual rate of reaction (Schaich, 2005; Sun et al., 2011). Treatment of barley seeds with H_2_O_2_ promotes transcription of GA biosynthesis genes and ABA catabolism genes (Ishibashi et al., 2017). The rate of O^2-^ produced by the mitochondria is proportional to the rate of oxygen consumption and local oxygen concentration in the tissue (Skulachev, 1996; Murphy, 2009; Møller, 2001). If HvAlaAT1 results in a relatively higher oxygen percentage in the embryo of non-dormant seeds during seed maturation, greater ROS accumulation due to autooxidation may lower the amount of oxygen consumption required to induce germination, resulting in a faster germination rate at 5% oxygen for non-dormant relative to dormant seeds. Nevertheless, it is unclear if the dormancy-associated alleles of SD1 significantly influence the enzymatic activity of HvAlaAT1, or if other aspects of the protein are influenced such as protein-protein interactions, stability or localization. Further work is needed to understand how this enzyme contributes to dormancy and how hypoxia influences dormancy more broadly. Monocot and dicot seed germination is influenced by hypoxia and the oxygen environment influences after-ripening and seed senescence (Bradford et al., 2008; Gerna et al., 2022; Bailly, 2023, Corbineau, 2022). This suggests that hypoxic responses are integrated into seed function across multiple plant species and thus identification of genetic loci that influence hypoxic response may provide novel targets for improving dormancy regulation more broadly. Altogether, this work provides insight into the genetic diversity of dormancy regulation in barley and highlights hypoxia responses as a key physiological process involved in this regulation.

## Supplementary data

**Supplementary Table S1:** Characteristics of the 215 lines tested from the ICARDA AM-14 population (Amezrou et al., 2018).

| AM-14 ID | Source | Adaptation | Germplasm | Row type |
| --- | --- | --- | --- | --- |
| AM-1 | ICARDA | Low Input | Breeding lines | 6 |
| AM-2 | ICARDA | Low Input | Breeding lines | 6 |
| AM-3 | ICARDA | Low Input | Breeding lines | 6 |
| AM-4 | ICARDA | Low Input | Breeding lines | 6 |
| AM-5 | ICARDA | Low Input | Breeding lines | 6 |
| AM-6 | ICARDA | Low Input | Breeding lines | 6 |
| AM-7 | ICARDA | Low Input | Breeding lines | 2 |
| AM-8 | ICARDA | Low Input | Breeding lines | 6 |
| AM-9 | ICARDA | Low Input | Breeding lines | 6 |
| AM-10 | ICARDA | Low Input | Breeding lines | 6 |
| AM-11 | ICARDA | Low Input | Variety | 2 |
| AM-12 | ICARDA | Low Input | Breeding lines | 2 |
| AM-13 | ICARDA | Low Input | Breeding lines | 6 |
| AM-15 | ICARDA | Low Input | Breeding lines | 6 |
| AM-16 | ICARDA | Low Input | Breeding lines | 6 |
| AM-18 | ICARDA | Low Input | Breeding lines | 6 |
| AM-19 | ICARDA | Low Input | Breeding lines | 6 |
| AM-20 | ICARDA | Low Input | Breeding lines | 6 |
| AM-21 | ICARDA | Low Input | Breeding lines | 6 |
| AM-22 | ICARDA | Low Input | Breeding lines | 6 |
| AM-23 | ICARDA | Low Input | Breeding lines | 6 |
| AM-24 | ICARDA | Low Input | Breeding lines | 6 |
| AM-25 | ICARDA | Low Input | Breeding lines | 6 |
| AM-26 | ICARDA | Low Input | Breeding lines | 6 |
| AM-27 | ICARDA | High Input | Breeding lines | 2 |
| AM-28 | ICARDA | High Input | Breeding lines | 6 |
| AM-29 | ICARDA | High Input | Breeding lines | 6 |
| AM-30 | ICARDA | High Input | Breeding lines | 2 |
| AM-31 | ICARDA | High Input | Breeding lines | 6 |
| AM-32 | ICARDA | High Input | Breeding lines | 2 |
| AM-33 | ICARDA | High Input | Breeding lines | 6 |
| AM-34 | ICARDA | High Input | Breeding lines | 6 |
| AM-35 | ICARDA | High Input | Breeding lines | 6 |
| AM-36 | ICARDA | High Input | Breeding lines | 6 |
| AM-37 | ICARDA | High Input | Breeding lines | 6 |
| AM-39 | ICARDA | High Input | Variety | 6 |
| AM-40 | ICARDA | High Input | Breeding lines | 2 |
| AM-41 | ICARDA | High Input | Variety | 6 |
| AM-42 | ICARDA | Low Input | Breeding lines | 6 |
| AM-43 | ICARDA | Low Input | Breeding lines | 2 |
| AM-44 | ICARDA | Low Input | Breeding lines | 2 |
| AM-45 | ICARDA | Low Input | Breeding lines | 6 |
| AM-46 | ICARDA | Low Input | Breeding lines | 2 |
| AM-47 | ICARDA | Low Input | Breeding lines | 2 |
| AM-48 | ICARDA | Low Input | Breeding lines | 2 |
| AM-49 | ICARDA | Low Input | Breeding lines | 6 |
| AM-50 | ICARDA | Low Input | Breeding lines | 2 |
| AM-51 | ICARDA | Low Input | Breeding lines | 2 |
| AM-52 | ICARDA | Low Input | Breeding lines | 2 |
| AM-53 | ICARDA | Low Input | Breeding lines | 6 |
| AM-54 | ICARDA | High input | Breeding lines | 2 |
| AM-55 | ICARDA | Low Input | Breeding lines | 2 |
| AM-56 | ICARDA | Low Input | Breeding lines | 2 |
| AM-57 | ICARDA | High Input | Breeding lines | 2 |
| AM-58 | ICARDA | Low Input | Breeding lines | 2 |
| AM-59 | ICARDA | High Input | Breeding lines | 2 |
| AM-60 | ICARDA | Low Input | Breeding lines | 6 |
| AM-62 | ICARDA | Low Input | Breeding lines | 6 |
| AM-63 | ICARDA | High Input | Breeding lines | 2 |
| AM-64 | ICARDA | Low Input | Breeding lines | 2 |
| AM-65 | ICARDA | Low Input | Breeding lines | 6 |
| AM-66 | ICARDA | High Input | Breeding lines | 6 |
| AM-67 | India | High Input | Variety | 6 |
| AM-68 | Syria | Low Input | Variety | 6 |
| AM-70 | ICARDA | High Input | Collection | 2 |
| AM-71 | ICARDA | Low Input | Breeding lines | 2 |
| AM-72 | ICARDA | Low Input | Breeding lines | 2 |
| AM-73 | ICARDA | Low Input | Breeding lines | 6 |
| AM-74 | ICARDA | Low Input | Breeding lines | 2 |
| AM-75 | ICARDA | Low Input | Breeding lines | 6 |
| AM-76 | ICARDA | Low Input | Breeding lines | 2 |
| AM-77 | ICARDA | Low Input | Breeding lines | 2 |
| AM-79 | ICARDA | Low Input | Breeding lines | 2 |
| AM-80 | ICARDA | Low Input | Breeding lines | 2 |
| AM-81 | ICARDA | Low Input | Breeding lines | 2 |
| AM-82 | ICARDA | Low Input | Breeding lines | 2 |
| AM-83 | ICARDA | Low Input | Breeding lines | 6 |
| AM-84 | ICARDA | Low Input | Breeding lines | 2 |
| AM-85 | ICARDA | Low Input | Breeding lines | 2 |
| AM-86 | ICARDA | Low Input | Breeding lines | 2 |
| AM-87 | ICARDA | Low Input | Breeding lines | 6 |
| AM-88 | ICARDA | Low Input | Breeding lines | 6 |
| AM-89 | ICARDA | Low Input | Breeding lines | 2 |
| AM-90 | ICARDA | Low Input | Breeding lines | 2 |
| AM-91 | ICARDA | Low Input | Variety | 6 |
| AM-92 | Morocco | Low Input | Variety | 6 |
| AM-93 | ICARDA | Low Input | Breeding lines | 6 |
| AM-94 | ICARDA | Low Input | Breeding lines | 2 |
| AM-95 | ICARDA | Low Input | Breeding lines | 6 |
| AM-96 | ICARDA | Low Input | Breeding lines | 6 |
| AM-97 | ICARDA | Low Input | Breeding lines | 6 |
| AM-98 | ICARDA | Low Input | Breeding lines | 2 |
| AM-99 | ICARDA | Low Input | Breeding lines | 2 |
| AM-100 | ICARDA | Low Input | Breeding lines | 6 |
| AM-101 | ICARDA | Low Input | Breeding lines | 2 |
| AM-102 | ICARDA | Low Input | Breeding lines | 2 |
| AM-103 | ICARDA | Low Input | Breeding lines | 2 |
| AM-104 | ICARDA | Low Input | Breeding lines | 2 |
| AM-105 | ICARDA | High Input | Breeding lines | 6 |
| AM-106 | ICARDA | Low Input | Breeding lines | 2 |
| AM-107 | ICARDA | Low Input | Breeding lines | 2 |
| AM-108 | ICARDA | High Input | Breeding lines | 6 |
| AM-109 | ICARDA | Low Input | Breeding lines | 2 |
| AM-110 | ICARDA | High input | Variety | 6 |
| AM-221 | ICARDA | High Input | Breeding lines | 6 |
| AM-222 | ICARDA | Low Input | Breeding lines | 6 |
| AM-223 | ICARDA | Low Input | Breeding lines | 6 |
| AM-224 | ICARDA | High Input | Breeding lines | 6 |
| AM-225 | ICARDA | Low Input | Breeding lines | 2 |
| AM-226 | ICARDA | Low Input | Breeding lines | 6 |
| AM-227 | ICARDA | High Input | Breeding lines | 6 |
| AM-228 | ICARDA | High Input | Breeding lines | 2 |
| AM-229 | ICARDA | High Input | Breeding lines | 2 |
| AM-230 | ICARDA | Low Input | Variety | 6 |
| AM-231 | Australia | Low Input | Variety | 2 |
| AM-232 | ICARDA | Low Input | Variety | 2 |
| AM-233 | ICARDA | Low Input | Variety | 2 |
| AM-234 | ICARDA | Low Input | Variety | 6 |
| AM-235 | ICARDA | Low Input | Variety | 6 |
| AM-236 | ICARDA | High Input | Breeding lines | 6 |
| AM-237 | ICARDA | High Input | Breeding lines | 6 |
| AM-238 | ICARDA | High Input | Breeding lines | 6 |
| AM-239 | USA | High Input | Variety | 6 |
| AM-240 | ICARDA | High Input | Breeding lines | 6 |
| AM-241 | Brazil | High Input | Variety | 6 |
| AM-242 | ICARDA | High Input | Variety | 6 |
| AM-243 | ICARDA | High Input | Breeding lines | 6 |
| AM-244 | ICARDA | High Input | Breeding lines | 6 |
| AM-245 | ICARDA | High Input | Breeding lines | 6 |
| AM-246 | ICARDA | High Input | Breeding lines | 6 |
| AM-247 | ICARDA | High Input | Breeding lines | 6 |
| AM-248 | ICARDA | High Input | Breeding lines | 6 |
| AM-249 | ICARDA | High Input | Breeding lines | 6 |
| AM-250 | ICARDA | High Input | Breeding lines | 6 |
| AM-252 | ICARDA | High Input | Breeding lines | 6 |
| AM-253 | India | Landrace | Variety | 6 |
| AM-254 | India | Landrace | Variety | 6 |
| AM-255 | India | Landrace | Variety | 6 |
| AM-256 | India | Landrace | Variety | 6 |
| AM-257 | India | High Input | Variety | 6 |
| AM-258 | India | Low Input | Variety | 6 |
| AM-261 | India | High Input | Variety | 6 |
| AM-262 | India | Low Input | Variety | 6 |
| AM-263 | India | High Input | Variety | 6 |
| AM-264 | India | High Input | Variety | 6 |
| AM-265 | India | High Input | Variety | 6 |
| AM-266 | India | High Input | Variety | 6 |
| AM-267 | India | Low Input | Variety | 6 |
| AM-268 | India | High Input | Variety | 2 |
| AM-269 | India | High Input | Variety | 2 |
| AM-270 | India | High Input | Variety | 2 |
| AM-271 | ICARDA | Low Input | Breeding lines | 2 |
| AM-272 | ICARDA | Low Input | Breeding lines | 2 |
| AM-273 | ICARDA | Low Input | Breeding lines | 2 |
| AM-274 | ICARDA | Low Input | Breeding lines | 2 |
| AM-277 | ICARDA | Low Input | Breeding lines | 2 |
| AM-278 | ICARDA | Low Input | Breeding lines | 2 |
| AM-279 | ICARDA | Low Input | Breeding lines | 2 |
| AM-280 | ICARDA | Low Input | Breeding lines | 2 |
| AM-281 | ICARDA | Low Input | Breeding lines | 2 |
| AM-282 | ICARDA | Low Input | Breeding lines | 2 |
| AM-283 | ICARDA | Low Input | Breeding lines | 6 |
| AM-284 | ICARDA | Landrace | Collection | 6 |
| AM-285 | ICARDA | Landrace | Collection | 6 |
| AM-286 | ICARDA | Landrace | Collection | 6 |
| AM-287 | ICARDA | Landrace | Collection | 6 |
| AM-288 | ICARDA | Landrace | Collection | 2 |
| AM-289 | ICARDA | Landrace | Collection | 6 |
| AM-290 | ICARDA | Landrace | Collection | 6 |
| AM-291 | ICARDA | Landrace | Collection | 6 |
| AM-292 | ICARDA | Landrace | Collection | 6 |
| AM-293 | ICARDA | Landrace | Collection | 6 |
| AM-294 | ICARDA | Landrace | Collection | 6 |
| AM-295 | ICARDA | Landrace | Collection | 6 |
| AM-296 | Australia | High input | Collection | 2 |
| AM-297 | Australia | High input | Variety | 2 |
| AM-298 | USA | Landrace | Collection | 2 |
| AM-299 | Australia | Low input | Variety | 2 |
| AM-300 | Australia | High input | Variety | 6 |
| AM-301 | Australia | High input | Variety | 6 |
| AM-302 | Australia | High input | Collection | 2 |
| AM-303 | Australia | High Input | Variety | 2 |
| AM-304 | USA | Landrace | Collection | 2 |
| AM-305 | USA | Landrace | Collection | 2 |
| AM-306 | USA | High Input | Variety | 6 |
| AM-307 | Australia | High input | Variety | 6 |
| AM-308 | Australia | High input | Variety | 6 |
| AM-309 | Australia | Low Input | Collection | 2 |
| AM-310 | USA | Landrace | Collection | 6 |
| AM-311 | Australia | High input | Variety | 2 |
| AM-312 | Australia | High input | Variety | 6 |
| AM-313 | Australia | High Input | Variety | 2 |
| AM-314 | China | High input | Variety | 2 |
| AM-315 | Australia | High Input | Variety | 2 |
| AM-316 | Australia | High Input | Variety | 2 |
| AM-317 | Australia | High Input | Variety | 2 |
| AM-318 | Australia | High Input | Variety | 2 |
| AM-319 | Australia | High input | Variety | 2 |
| AM-320 | Australia | High input | Collection | 2 |
| AM-321 | ICARDA | Low Input | Collection | 6 |
| AM-322 | USA | Landrace | Collection | 6 |
| AM-323 | Australia | High input | Collection | 6 |
| AM-324 | Australia | Landrace | Collection | 6 |
| AM-325 | Japan | Landrace | Variety | 6 |
| AM-326 | ICARDA | Landrace | Collection | 6 |
| AM-327 | Australia | High input | Variety | 2 |
| AM-328 | Australia | High input | Variety | 6 |
| AM-329 | ICARDA | Landrace | Collection | 6 |
| AM-330 | ICARDA | Low Input | Breeding lines | 6 |
| AM-331 | ICARDA | Low Input | Breeding lines | 6 |
| AM-332 | ICARDA | Low Input | Breeding lines | 6 |
| AM-333 | ICARDA | Low Input | Breeding lines | 6 |
| AM-334 | ICARDA | Low Input | Breeding lines | 6 |
| AM-335 | ICARDA | Low Input | Breeding lines | 6 |
| AM-336 | ICARDA | Low Input | Breeding lines | 2 |

**Supplementary Table S2:**
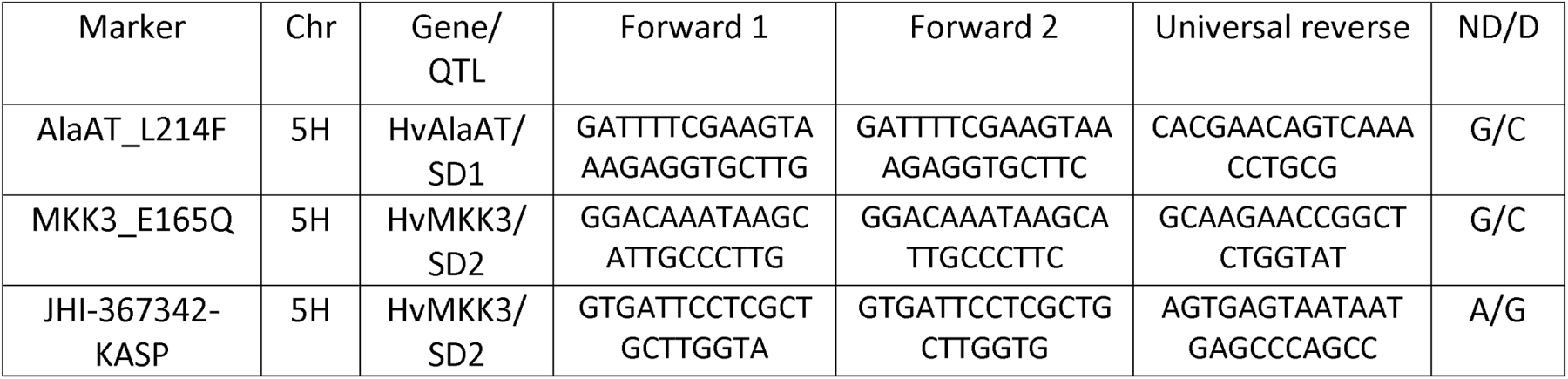
KASP primers used for determination of SD1 and SD2 alleles (Sweeney et al., 2022).

**Supplementary Table S3:**
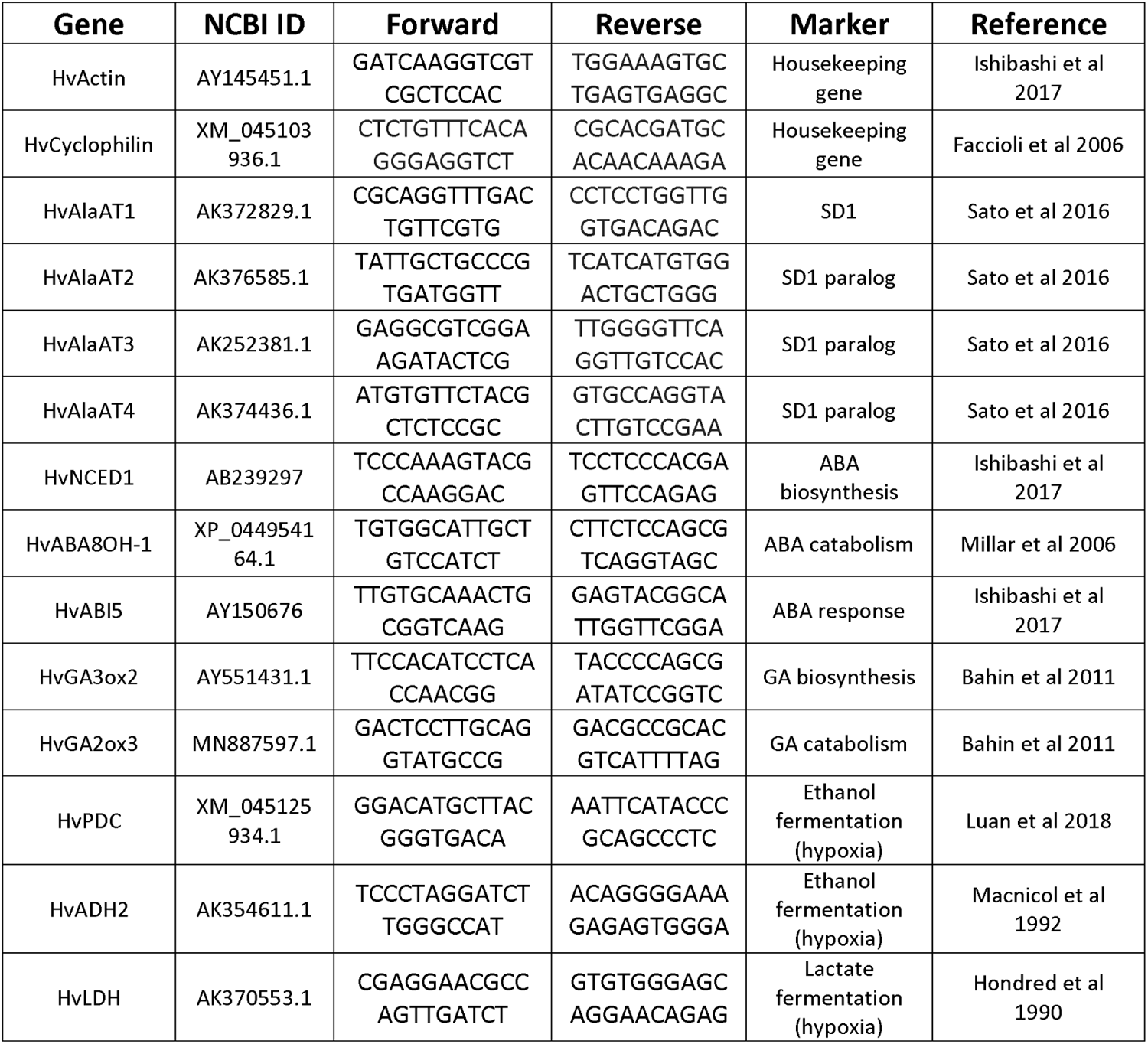
Primers for qPCR during grain maturation of barley heads from the LegCi population.

**Supplementary Table S4:** Gene expression during grain maturation in select lines from the LegCi population. Barley spikes were harvested 15 days after anthesis from 4 independent plants per genotype within an experiment. The average expression (± SD) of the indicated gene from a single experiment was normalized to actin and cyclophilin expression and expressed relative to the parental lines Leger or CI9831. No significant differences between progeny lines relative to their most genetically similar parental line was observed (H106-311, ND vs CI9831, D)(H106-374, D vs Leger, ND) using t-tests (p<0.05). The experiment was repeated three times with similar results.

| Gene | Inducer | Barley genotype |  |  |  |  |  |  |  |
| --- | --- | --- | --- | --- | --- | --- | --- | --- | --- |
|  |  | CI9831 |  | H106-311 |  | Leger |  | H106-374 |  |
|  |  | Avg | SD | Avg | SD | Avg | SD | Avg | SD |
| *AlaAT1* | | 1.2 ± | 1.13 | 0.3 ± | 0.15 | 1.1 ± | 0.49 | 1.5 ± | 0.55 |
| *AlaAT2* | | 1.0 ± | 0.35 | 0.7 ± | 0.28 | 1.0 ± | 0.22 | 0.9 ± | 0.40 |
| *AlaAT3* | | 1.0 ± | 0.33 | 0.9 ± | 0.46 | 1.0 ± | 0.37 | 0.8 ± | 0.42 |
| *AlaAT4* | | 1.0 ± | 0.52 | 1.0 ± | 0.66 | 1.0 ± | 0.26 | 0.7 ± | 0.45 |
| *AlaAT5* | | 1.0 ± | 0.19 | 1.1 ± | 0.25 | 1.0 ± | 0.16 | 0.9 ± | 0.28 |
| *GA3ox2* | GA | 0.9 ± | 0.22 | 0.9 ± | 0.81 | 2.0 ± | 0.56 | 6.9 ± | 7.66 |
| *GA2ox3* | GA | 1.0 ± | 0.39 | 0.3 ± | 0.12 | 1.0 ± | 0.42 | 0.8 ± | 0.60 |
| *ABA8H-1* | ABA | 1.0 ± | 0.42 | 0.3 ± | 0.15 | 1.2 ± | 1.00 | 1.0 ± | 0.74 |
| *ABI5* | ABA | 1.1 ± | 0.57 | 0.3 ± | 0.22 | 1.0 ± | 0.11 | 0.8 ± | 0.41 |
| *NCED1* | ABA | 1.4 ± | 1.34 | 0.1 ± | 0.06 | 1.0 ± | 0.38 | 1.1 ± | 0.40 |
| *LDH* | Hypoxia | 0.9 ± | 0.24 | 0.8 ± | 0.25 | 1.1 ± | 0.50 | 1.9 ± | 1.06 |
| *PDC* | Hypoxia | 1.1 ± | 0.62 | 0.3 ± | 0.24 | 1.1 ± | 0.56 | 1.5 ± | 0.37 |

**Supplementary Fig. S1:**
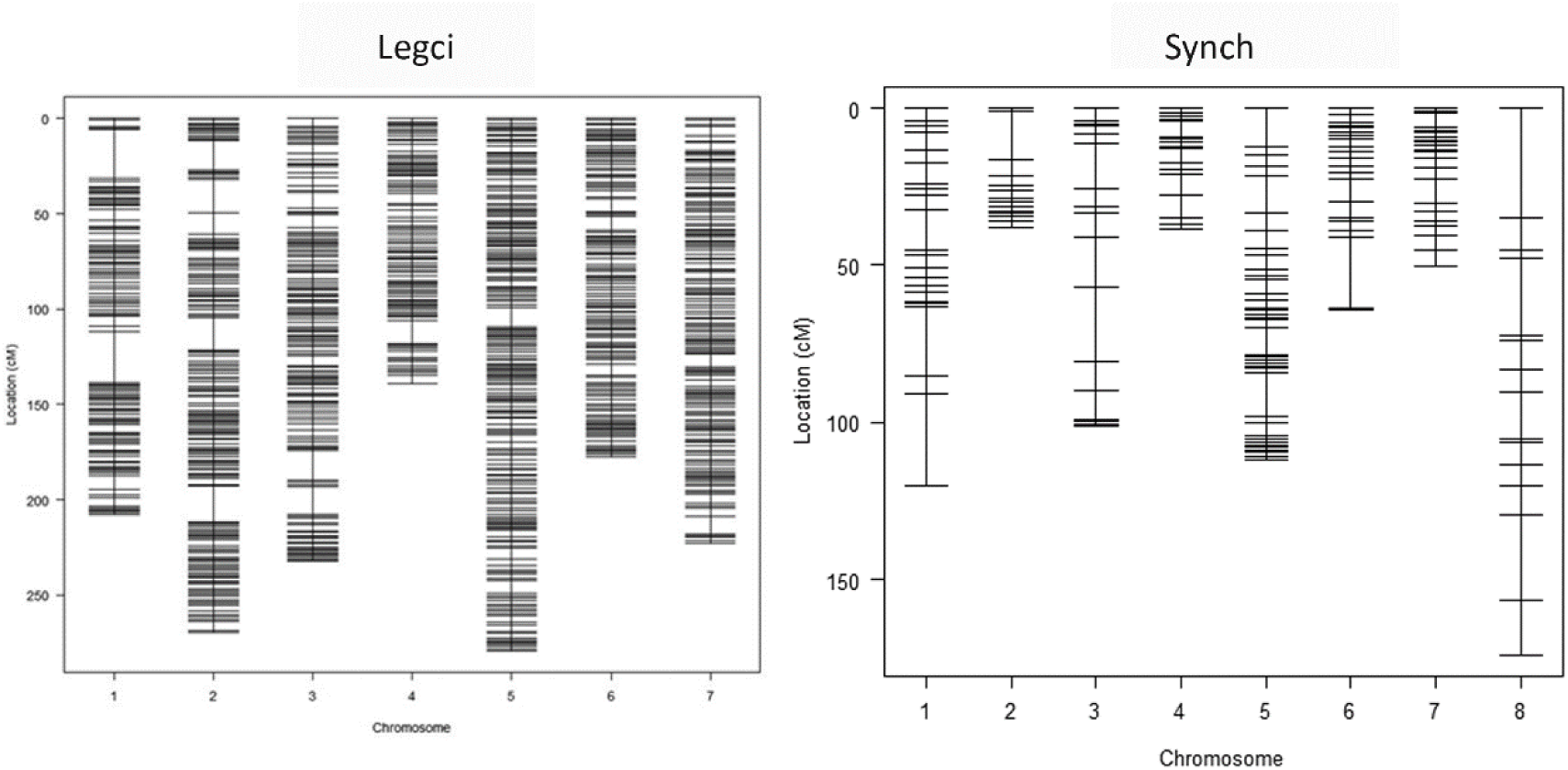
SynCH recombinant inbred line population linkage map based on 2827 markers. Linkage map constructed using IciMapping (Meng et al., 2015). Linkage map for the LegCi double haploid population based on 963 GBS markers (Abed et al., 2022). Linkage map was constructed using MSTmap (Wu et al., 2008).

**Supplementary Fig. S2:**
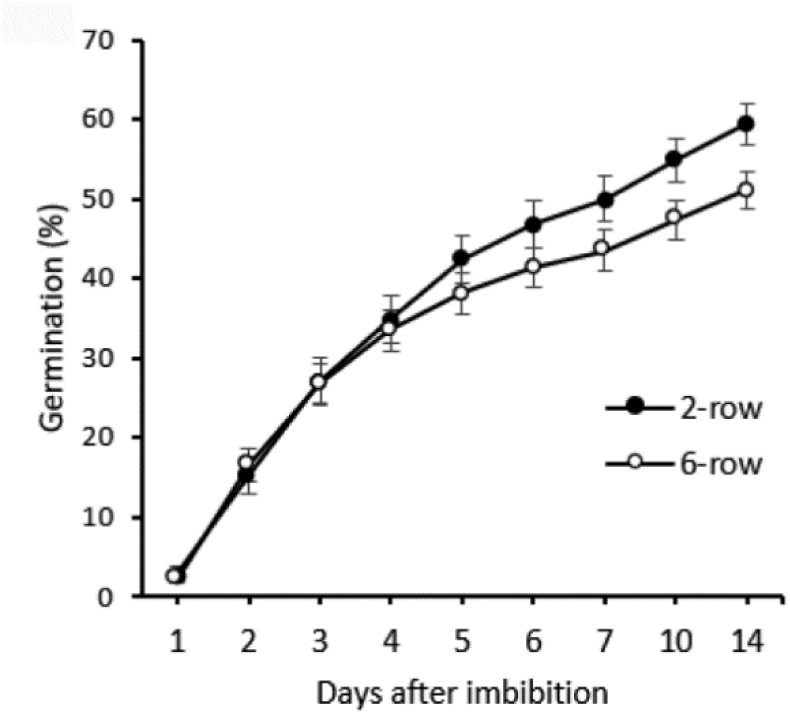
Germination percentage trends for the ICARDA AM-14 panel and relationship with phenotypic characteristics. For each genotype, 20 seeds were distributed in each of three replicate plates and germination was monitored at the indicated days after imbibition (DAI). The average germination percentage ± standard error for all timepoints separated by documented adaptation (118 low input, 21 landrace (not shown), 76 high input) is shown. No significant differences between groups was observed and the experiment was repeated once.

**Supplementary Fig. S3:**
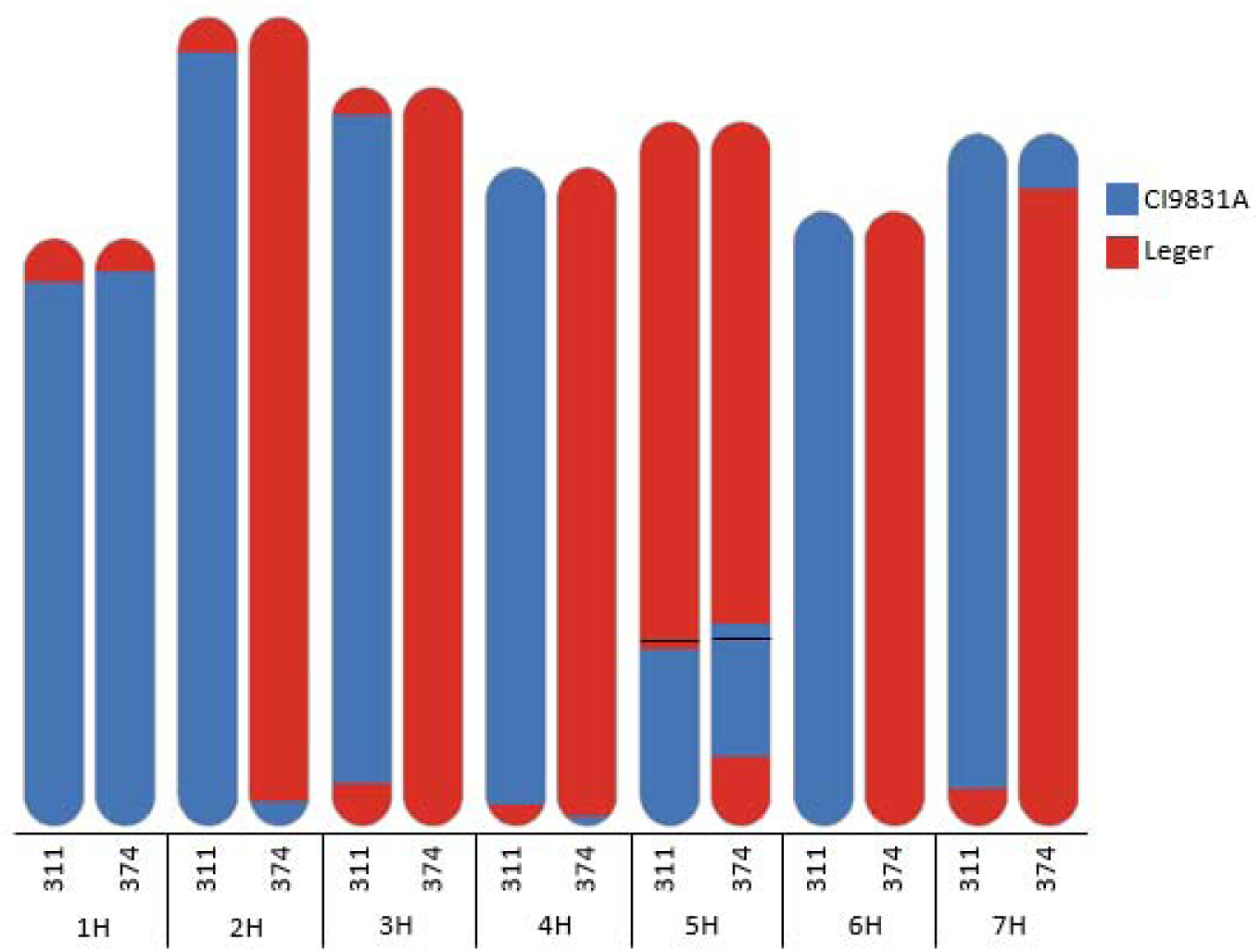
Alignment of shared molecular markers between doubled haploid recombinant lines H106-311 and H106-374 and their parental lines CI9831 (blue) or Leger (red). Relationship to parental lines was determined by 963 GBS markers. The solid black line on chromosome 5H indicates the approximate location of the HvAlaAT1/SD1 locus based on the physical location from the Morex v1 IBSCv2 reference genome (Mascher et al., 2017).

**Supplementary Fig. S4:**
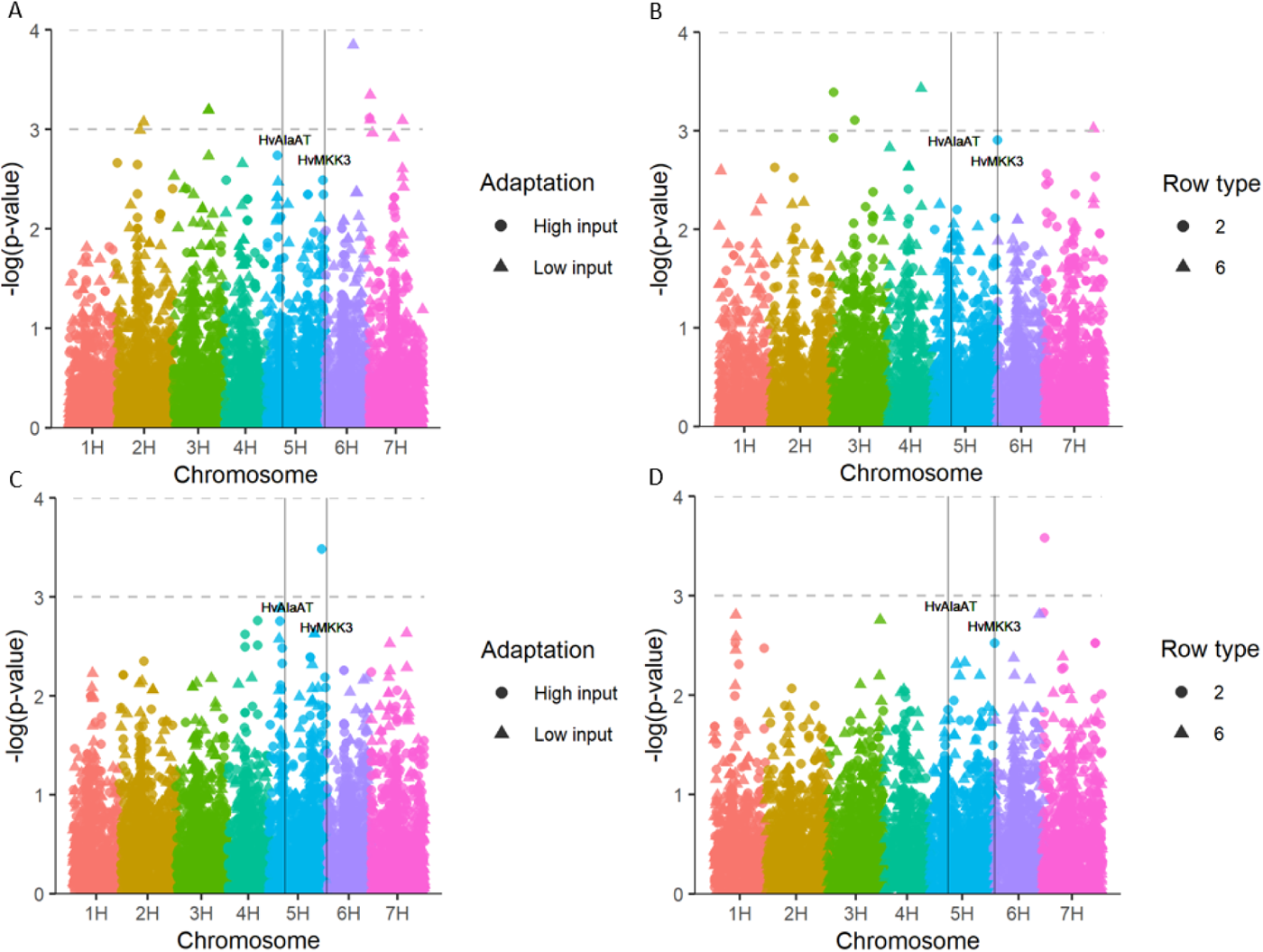
Association between (A, B) 2 DAI germination percentage or (C, D) 14 DAI with markers from AM-14 lines of (A, C) adaptation or (B, D) row type mapped independently. Approximate location of SD1 and SD2 was determined using the closest marker physical position to HvAlaAT1 and HVMKK3 in the Morex v3 reference genome (Mascher et al., 2021).

**Supplementary Fig. S5:**
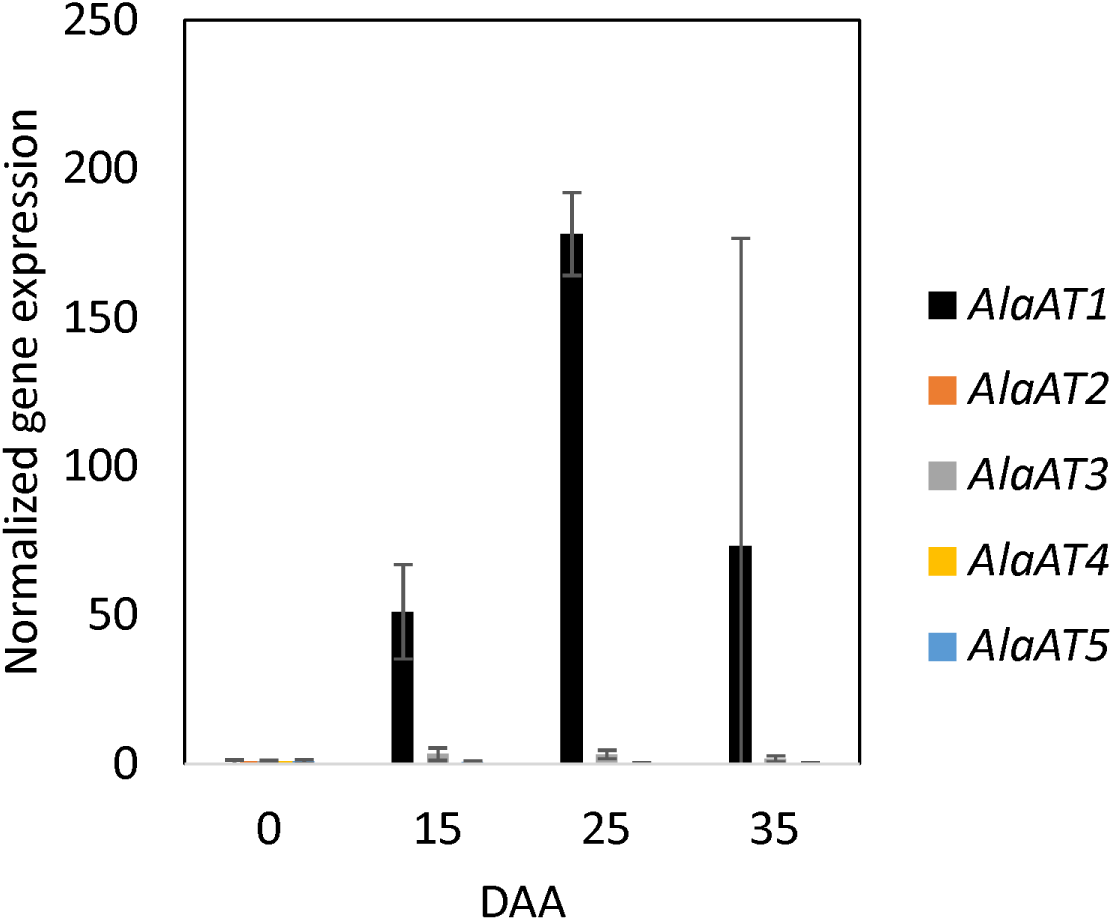
Alanine aminotransferase gene expression during grain maturation in Leger. Barley spikes were harvested at the indicated number of days after anthesis from 4 independent plants per genotype within an experiment. The average expression (± SD) of the indicated gene from a single experiment was normalized to actin and cyclophilin expression and expressed relative to 0 DAA, at anthesis.

**Supplementary Fig. S6:**
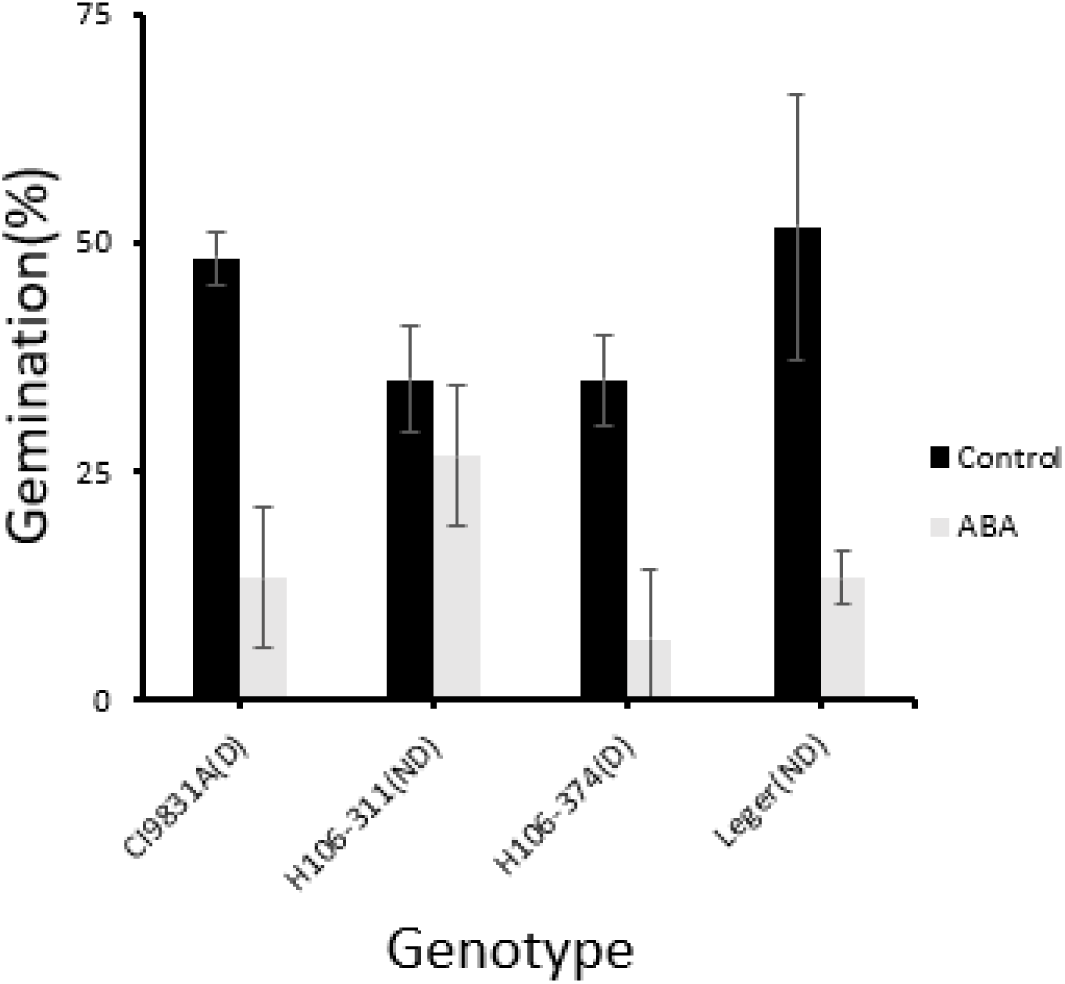
Effect of 20 µM ABA treatment on LegCi genotypes with either (D) dormant or (ND) non-dormant alleles at the HvAlaAT_L214F locus. 20 seeds germinated in darkness at 20°C sampled once every 24 hours 1-6 days after imbibition. Values represent average germination percentage ± standard deviation 5 days after imbibition (n=3). Seeds were scored as germinated at any sign of radical emergence to separate the influence ABA has on germination from its effect on post-germination growth.

**Supplementary Fig. S7:**
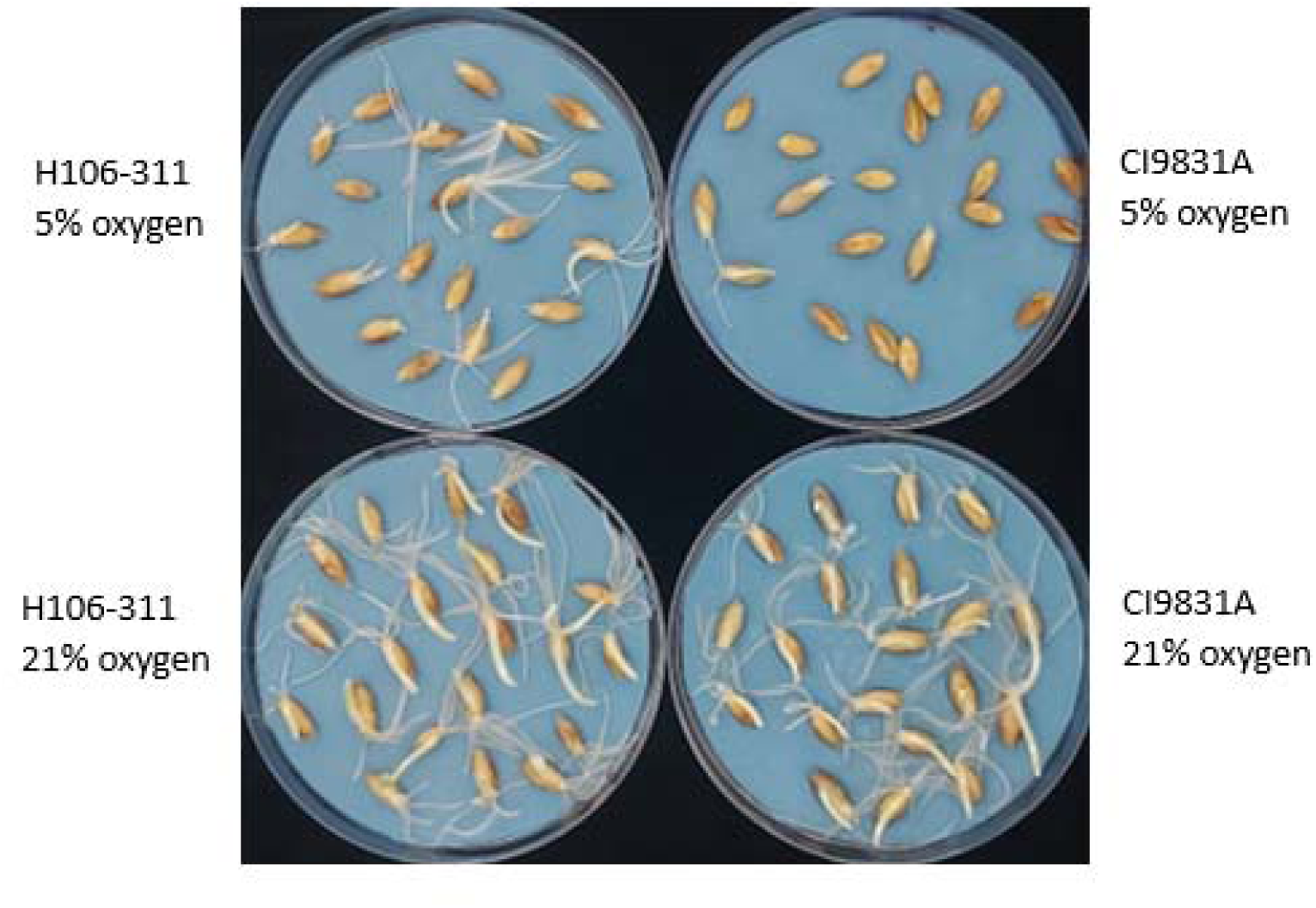
Representative photo of hypoxia sensitivity during germination in the (D) CI9831 and (ND) H106-311 comparison. Photo was taken three days after imbibition. All seeds are dehulled.

Supplemental Data 1: Number of germinated barley seeds after imbibition in three replicate plates (A-C) across three populations and heritability (H2) of germination % for each day.

## Acknowledgements

Thanks to the technical staff at Agriculture and Agri-Food Canada for training in KASP and hypoxia assays including Martin Charette, Ulrica McKim and Ryan Tobalt. Thanks also to the staff and students who helped to set up germination experiments including Julia Desbiens, Monique Power, Kirsten Holy and Kim Madge and to Monique Power for her feedback on the manuscript. Thanks to the greenhouse staff at the Ottawa Research and Development Center for maintaining the plants used for these experiments. This research was supported by Agriculture & Agri-Food Canada’s Genomics Research and Development Initiative.

## Author Contributions

L.F. and E.K.B. designed the experiments and analyzed the results. L.F. performed the experiments with the exception of one gene expression experiment, which was executed by E.K.B. W.B. performed the SynCH population analysis and assisted with GWAS and QTL analyses. B.S. and R.K. provided germplasm and genotyping data and advice on experimental set up. E.K.B. and L.F. co-wrote the manuscript with input from all co-authors.

